# IMPAIRED NEURAMINIDASE AND POLYMERASE ACTIVITIES CORRESPOND WITH LIMITED AEROSOL INFECTIVITY OF B3.13 AND D1.1 H5N1 LINEAGES IN HUMAN RESPIRATORY CULTURES

**DOI:** 10.64898/2026.08.18.745466

**Authors:** L. Claire Gay, Dikshya Regmi, Flavio Cargnin Faccin, Ignacio Scanarotti, Alison Mark, C. Joaquín Cáceres, Teresa Mejias, Maria Corkran, Margaret A. Scull, Rafael Medina, Adolfo Garcia-Sastre, Daniel R. Perez

## Abstract

The ongoing panzootic of clade 2.3.4.4b highly pathogenic avian influenza (HPAI) H5N1 viruses has reached a critical point, marked by unprecedented mammalian spillover and sustained outbreaks in U.S. dairy cattle. While these viruses remain highly lethal in traditional ferret models, human infections—primarily linked to the B3.13 and D1.1 lineages—have been notably mild, typically presenting as conjunctivitis with minimal respiratory involvement. In this study, we address this disconnect by evaluating the infectivity of recent H5N1 isolates using a physiologically relevant air-liquid interface (ALI) culture system that incorporates an aerosol settling chamber. We demonstrate that while direct liquid inoculation leads to efficient replication, aerosolized H5N1 strains exhibit a significant defect in their ability to infect human respiratory epithelium. In contrast, a prototypic H5N1 virus remains highly pathogenic and lethal in ferrets regardless of the inoculation route, showing systemic dissemination to the brain and other organs. Our findings identify two primary viral determinants driving this respiratory restriction: reduced neuraminidase (NA) enzymatic activity and impaired polymerase activity.

Collectively, these results suggest that commonly used mammalian models may overstate current human pandemic risk. This work highlights the critical need for alternative risk-assessment platforms to identify the specific genetic shifts required for these viruses to overcome existing barriers to human adaptation.

**IMPORTANCE:** Current pandemic risk assessments rely heavily on animal models, particularly ferrets, to evaluate the threat posed by emerging influenza viruses. However, recent H5N1 viruses that have caused predominantly mild human infections have remained highly virulent in these models, creating uncertainty about how well they predict human disease. Using a physiologically relevant human airway model infected through aerosol exposure, we demonstrate that contemporary H5N1 are markedly restricted in their ability to establish respiratory infection despite retaining high virulence in ferrets. These findings highlight the importance of incorporating human airway aerosol models into pandemic risk assessment frameworks to better evaluate the human adaptation and respiratory infection potential of emerging influenza viruses.

## INTRODUCTION

The evolutionary history of highly pathogenic avian influenza (HPAI) H5N1 is defined by the emergence of the A/goose/Guangdong/1/1996 (Gs/Gd) lineage, which has fundamentally altered the global landscape of influenza virus ecology and public health (1). Since its initial detection in domestic waterfowl in southern China, the Gs/Gd hemagglutinin (HA) gene has exhibited extraordinary plasticity, diversifying into ten primary ancestral clades (0–9) and hundreds of sub-clades (2). While early global expansions were driven by clades such as 2.2 (2005) and 2.3.2.1 (2008), the last decade has been dominated by the rapid diversification of clade 2.3.4.4 (3). This specific lineage is characterized by its high propensity for reassortment with local low-pathogenicity avian influenza (LPAI) viruses, leading to a "shuffling" of NA segments that has produced various H5Nx subtypes (including H5N2, H5N5, H5N6, and H5N8) (4).

The 2.3.4.4 lineage is unique for its sustained global circulation and its ability to thrive in a broad range of wild bird hosts, which act as the primary vectors for intercontinental transmission. Within this lineage, several distinct subclades have reached high regional or global significance, most notably clades 2.3.4.4a, b, c, and h. Clade 2.3.4.4h is primarily circulating in Asia (particularly China and Southeast Asia) and parts of Egypt, often associated with H5N6 and H5N1 subtypes in poultry and live bird markets (5). Clades 2.3.4.4a and 2.3.4.4c are historically significant for the first major H5N8 waves in 2014–2016 across Eurasia and North America, though they have largely been displaced by the "b" subclade. Clade 2.3.4.4b is currently the most dominant and geographically widespread lineage globally (6). Since late 2020, it has caused an unprecedented panzootic, affecting over 500 bird species and more than 80 mammalian species (7).

The arrival of clade 2.3.4.4b in the Western Hemisphere marked a critical turning point in the panzootic. In December 2021, the virus was first detected in Newfoundland, Canada, likely introduced via the North Atlantic flyway by migratory birds from Europe. By early 2022, the virus had infiltrated all four North American migratory flyways, causing catastrophic losses in commercial poultry and severe mortality in wild raptors and scavengers (6).

The virus moved south and entered Central and South America in late 2022, reaching the Neotropical region for the first time in recorded history (8). This southward expansion led to massive mortality events in colonial seabirds and pinnipeds (seals and sea lions) along the Pacific and Atlantic coasts, particularly in Peru, Chile, and Argentina (8–10). The 2.3.4.4b lineage in the Americas has further diverged into critical genotypes through reassortment with North American wild bird viruses. These include the B3.2 genotype responsible for the initial massive spread and mammalian mortality in South America and the subsequent introduction into Antarctica (10, 11). In March 2024, H5N1 HPAI of the clade 2.3.4.4b lineage was detected in dairy cattle (12). Infected animals exhibited clinical signs including nasal discharge, lethargy, and profound reductions in milk production (13, 14). Although H5N1 viruses typically infect the respiratory tract, in dairy cattle the virus was predominantly detected in mammary tissue and milk (13). Several human cases were subsequently reported, primarily among dairy workers, and were generally associated with conjunctivitis (15). Nevertheless, cases of severe human disease, including fatal infections, from the recent clade 2.3.4.4b H5N1 viruses have been reported (16). Epidemiological investigations suggested transmission was likely facilitated by direct contact during milking (17). The D1.1 genotype represents a recent reassortant (late 2024/2025) that has shown increased fitness in wild birds and was linked to the first H5N1 fatalities in the Americas (Louisiana, USA and Durango, Mexico) (18). Overall, reported mortality associated with 2.3.4.4b clade viruses appears reduced compared to earlier H5N1 strains (19, 20), with a global human case fatality rate (CFR) of approximately 14% reported between December 2023 and December 2025, compared with an overall CFR of 48% from January 2003 to December 2025 (19, 20).

Despite the relatively mild clinical presentation observed in most human infections, clade 2.3.4.4b HPAI viruses cause severe disease and high mortality in mammalian models of influenza commonly used for human risk inference (21–26). Multiple studies have evaluated the ability of these viruses to transmit via direct contact or aerosol respiratory contact (21, 24, 26–31). Depending on the viral strain, aerosol contact transmission was observed; however, lethality was consistently demonstrated in the ferret model (21, 24–28, 30, 31). Notably, most pathogenesis studies in these models rely on a relatively large liquid inoculation dose (100-1000 µL), by either or both the intranasal or ocular routes, but not through airborne exposure via aerosols, except for a single study evaluating low-dose aerosol exposure that reported lethal infection of clade 2.3.4.4b H5N1 in ferrets (26).

In parallel, several studies have assessed the ability of clade 2.3.4.4b H5N1 viruses to replicate in human respiratory air-liquid interface (ALI) cultures (22, 25, 26, 32). Collectively, these studies demonstrate efficient viral replication across multiple ALI culture systems and multiplicities of infection (MOI) when viruses are introduced via liquid inoculation. To our knowledge, aerosol inoculation of ALI cultures with clade 2.3.4.4b H5N1 viruses has not been reported. Thus, while human infections with clade 2.3.4.4b viruses are generally associated with mild disease and low mortality (15, 19, 20), experimental studies in mammalian models frequently demonstrated high viral replication and lethality, underscoring a disconnect between animal model-based and human disease outcomes and highlighting the need for improved approaches to assess pandemic risk.

In this study, we compare direct liquid and aerosol (nebulizer) inoculation *in vitro* to define the impact of inoculation route on the pathogenesis and replication dynamics of clade 2.3.4.4b HPAI viruses, specifically of those derived from the B3.13, D1.1, and (ancestral) A1 lineages. Human isolates were selected to facilitate direct comparison between observed human disease and experimental outcomes in ALI systems. Our findings indicate a restrictive ability of 2.3.4.4b H5N1 viruses to infect human respiratory ALI cultures by the airborne route, due in part to reduced NA activity as well as impaired polymerase activity. Our observations in the ALI system are consistent with the limited involvement of the respiratory tract in human cases of 2.3.4.4b H5N1 infections and suggest that commonly used mammalian models may overestimate human susceptibility and disease severity. Together, these results emphasize the need for alternative physiologically relevant experimental systems to accurately assess the pathogenesis and pandemic risk of emerging influenza viruses.

## RESULTS

### Decreased susceptibility of HAE/ALI cells to aerosolized H5N1 virus infection

In previous work, we established an aerosol settling chamber system to characterize the airborne infectivity of influenza A viruses (IAVs) using a physiologically relevant in vitro model (33). By utilizing immortalized, differentiated human airway epithelial (HAE clone BCi-NS1.1) (34) ALI cultured cells, this platform allows for aerosol infection of influenza viruses. Our evaluation of multiple IAV subtypes—including pandemic 2009 H1N1, seasonal swine H3N2, and avian H9N2—demonstrated that while liquid bolus delivery often results in uniform infectivity across strains, the aerosol system effectively differentiates their infectious potential based on the airborne infectious dose 50 (AID_50_) (33). Thus, this system serves as a scalable alternative to resource-intensive in vivo transmission studies. We expanded the use of this system to assess the susceptibility of our HAE/ALI cells to clade 2.3.4.4b H5N1 infection (Fig. 1a). Specifically, HAE/ALI cells in 12-well plates were washed and then simultaneously infected via direct inoculation (DI) or aerosol exposure (AR) at MOIs of 0.1 or 1. Transepithelial electrical resistance (TEER) was measured prior to each infection with HAE/ALI cultures with a subset of wells to assess viability of tight junctions in the cultures. Analysis of HAE/ALI culture integrity confirmed high TEER values (range, 350-1,300 Ω•cm^2^), indicative of strong, healthy barrier with tight intercellular junctions and low permeability prior to infection (Fig. 1b), consistent with previous work (34–36). Additionally, MDCK cells in 12-well plates placed in the chamber at the same time as the HAE/ALI cells were used as positive controls for virus infectivity. We analyzed the number of wells positive for infectious virus after aerosol exposure or direct inoculation and the amount of virus produced in each well over a 72 h period with samples collected at 24, 48, and 72 hours post-infection (hpi). The H5N1 viruses tested included A/Texas/37/24 clade 2.3.4.4b B3.13 (TX/24), A/British Columbia/PHL-2032/24 clade 2.3.4.4b D1.1 (BC/24), and A/turkey/Indiana/3707-003/22 (ty/IN/22) clade 2.3.4.4b A1. With the exception of ty/IN/22, which was a field isolate, the remaining viruses were generated by reverse genetics. All viruses carried avian-like adapted mutations with the exception of TX/24 which naturally carries the PB2 E627K mutation that favors replication in mammalian cells (22). In order to establish the role of the polybasic cleavage site of HA for infectivity, we additionally produced the A/Texas- Halo/37/24 (ΔTX/24) virus carrying the low pathogenic cleavage site derived from the H1 HA segment of the laboratory-adapted PR8 strain. The infectivity was compared to those established using the prototypic 2009 pandemic (mouse-adapted) A/California/04/2009 (H1N1) strain (ma-Ca04), which we have previously characterized (33).

**Figure 1.**
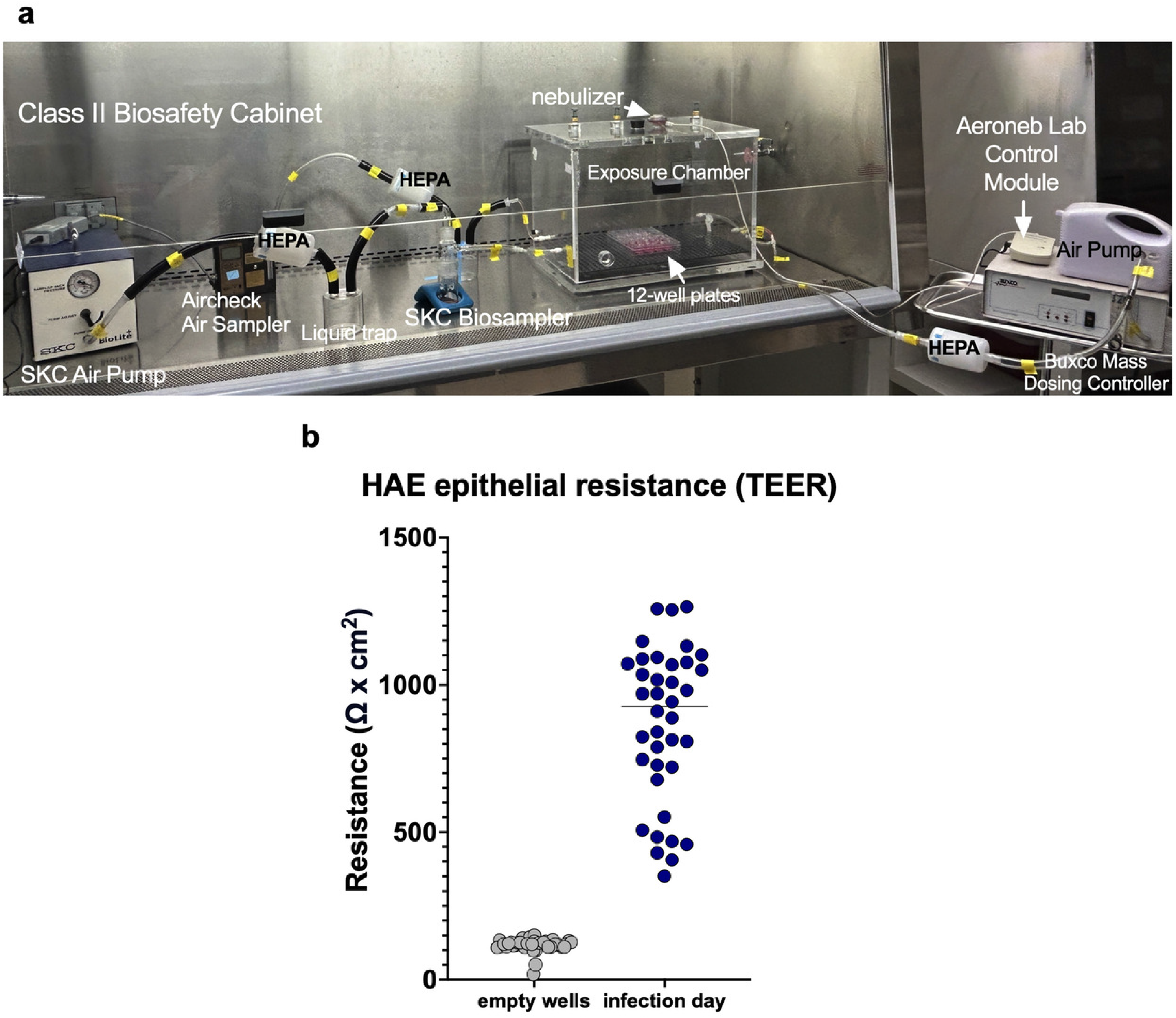
Aerosol exposure set up and TEER measurements for barrier integrity control. **(a)** Photograph depicting the experimental aerosol exposure system housed inside a Class II Biosafety Cabinet. The setup illustrates the central exposure chamber containing 12-well plates, an aerosol generation system (nebulizer and control modules), and the integrated air sampling and filtration circuit (HEPA filters, liquid trap, biosamplers, and air pumps). **(b)** Transepithelial electrical resistance (TEER) was measured prior to infection in a subset of HAE/ALI wells. Data points represent individual wells, with median values indicated. Statistical significance was determined using an Unpaired Welch’s t-test. Welch’s correction was preferred over the standard unpaired t test because the groups may have unequal variability.

At MOI 0.1, all H5N1 viruses tested showed decreased infectivity and replication when aerosolized for 15 min in the chamber (Fig. 2). At 24 hpi, TX/24 and ΔTX/24 had 3/12 and 2/12 wells positive, respectively, whereas both BC/24 and ty/IN/22 showed 0/12 positive wells. In contrast, the ma-Ca04 strain infected 12/12 wells (Fig. 2a). The limited infectivity of the H5N1 strains was also observed at 48 hpi (Fig. 2b) and persisted through 72 hpi (Fig. 2c), with TX/24 infecting 4/12, ΔTX/24 2/12 wells, and BC/24 and ty/IN/22 remaining undetectable. In contrast, ma-Ca04 maintained full infectivity, consistent with our previous studies (33). Consistent with active virus replication, viral titers increased over time in positive wells. In contrast to aerosol exposure, direct inoculation at MOI 0.1 showed higher infectivity in HAE cultures, in a strain-dependent manner. By 72 hpi, TX/24, ΔTX/24, and ma-Ca04 infected all wells, BC/24 infected 6/12 wells, whereas ty/IN/22 remained unable to infect HAE cells (Fig. 2a-c).

**Figure 2:**
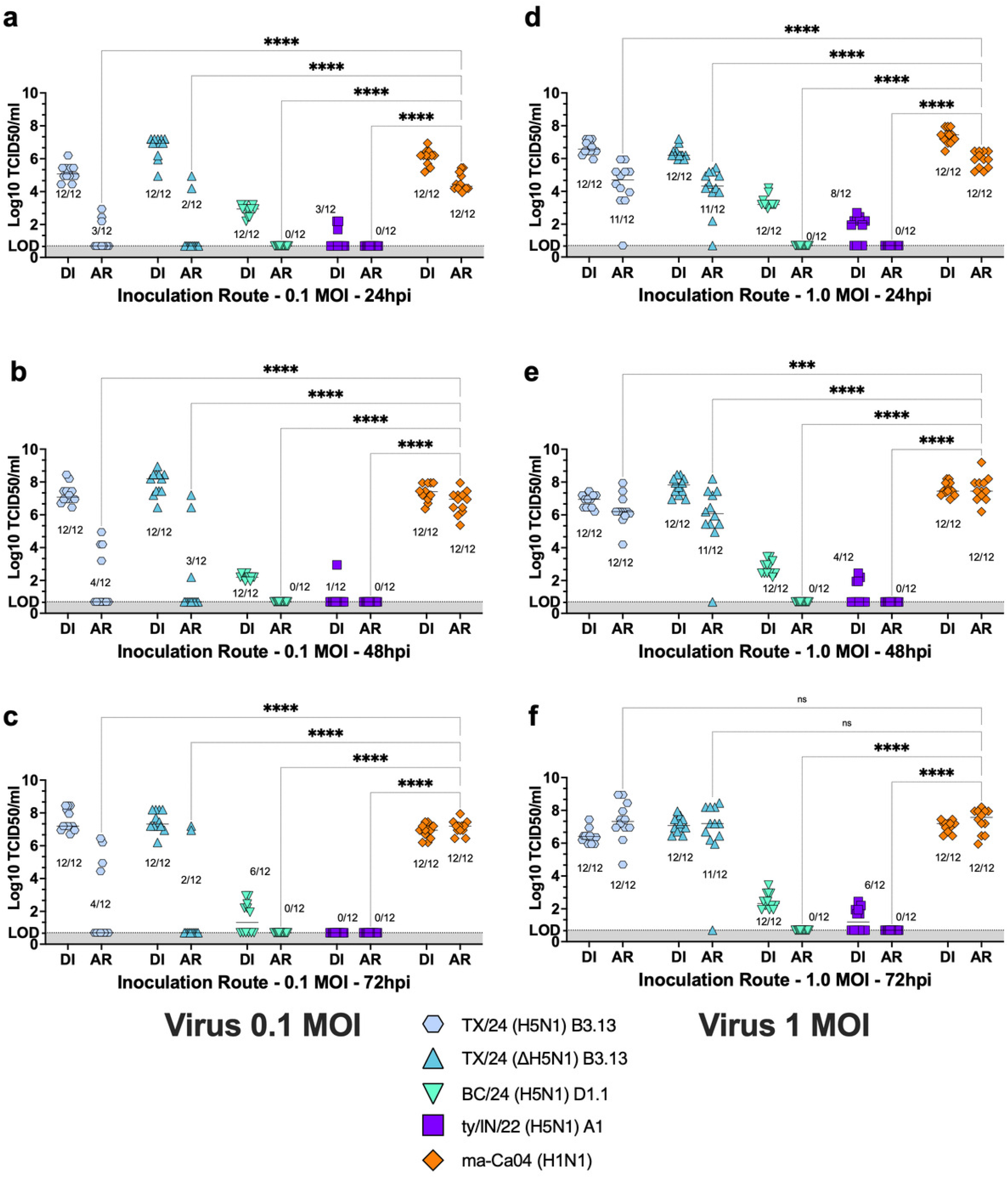
Reduced susceptibility of HAE cells to aerosolized H5N1 infection. HAE/ALI cultures were infected by direct inoculation (DI) or aerosol exposure (AR) at an MOI of 0.1 **(a-c)** or 1 **(d-f)** with A/Texas/37/24 (H5N1) (TX/24), A/Texas-Halo/37/24 (H5N1) (ΔTX/24), A/British Columbia/PHL-2032/24 (H5N1) (BC/24), A/turkey/Indiana/3707-003/22 (H5N1) (ty/IN/22), or A/California/04/2009 (H1N1) (ma-Ca04). Samples were collected at 24, 48, and 72 h post- infection, and viral titers were determined by TCID_50_ assay. Data are presented as individual wells with median values indicated. Statistical significance was determined using an ordinary two-way ANOVA followed by Dunnett’s multiple comparisons test. For visual clarity, only select significance levels are displayed on the graphs; comprehensive results are available in Supplementary Table 2. Asterisks denote statistical significance: ****P < 0.0001, ***P < 0.001, **P < 0.01, *P < 0.05; ns = not significant.

At an MOI of 1, aerosol exposure resulted in partial restoration of infectivity (Fig. 2d–f). By 72 hpi, TX/24 and ΔTX/24 reached 12/12 and 11/12 positive wells, respectively; conversely, BC/24 and ty/IN/22 remained undetectable across all time points. Direct inoculation at 1 MOI further enhanced infectivity for TX/24, ΔTX/24, BC/24, and ma-Ca04, with all reaching 12/12 positive wells. However, viral yields remained strain dependent. Despite reaching full well-positivity, BC/24 exhibited significantly lower titers compared with TX/24, ΔTX/24, and ma-Ca04, suggesting an inherent infectivity defect in HAE/ALI cells. Similarly, direct inoculation with ty/IN/22 yielded only 6/12 positive wells by 72 hpi, further highlighting the restricted infectivity of this strain in the HAE/ALI model.

To determine whether the aerosolization process was responsible for the reduced infectivity of H5N1 viruses in HAE/ALI cells, we titrated virus recovered from MDCK cells exposed to aerosols alongside the HAE/ALI cultures (Fig. 3). These data demonstrated that the H5N1 strains were able to maintain infectivity following aerosol delivery in MDCK cells. Specifically, TX/24 and ΔTX/24 exhibited robust infectivity comparable to the ma-Ca04 control, reaching 12/12 positive wells with similar viral titers. Although BC/24 and ty/IN/22 showed delayed growth kinetics, both reached 12/12 positive wells by 72 hpi in MDCK cells, with titers comparable to the other strains at an MOI of 1. Notably, ty/IN/22 yielded lower viral titers overall, regardless of the MOI or inoculation route, suggesting an inherent growth defect of this strain in MDCK cells (Fig. 3). Collectively, these results indicate that the previously observed infectivity defects are specific to the HAE/ALI system rather than a result of the aerosolization process itself.

**Figure 3:**
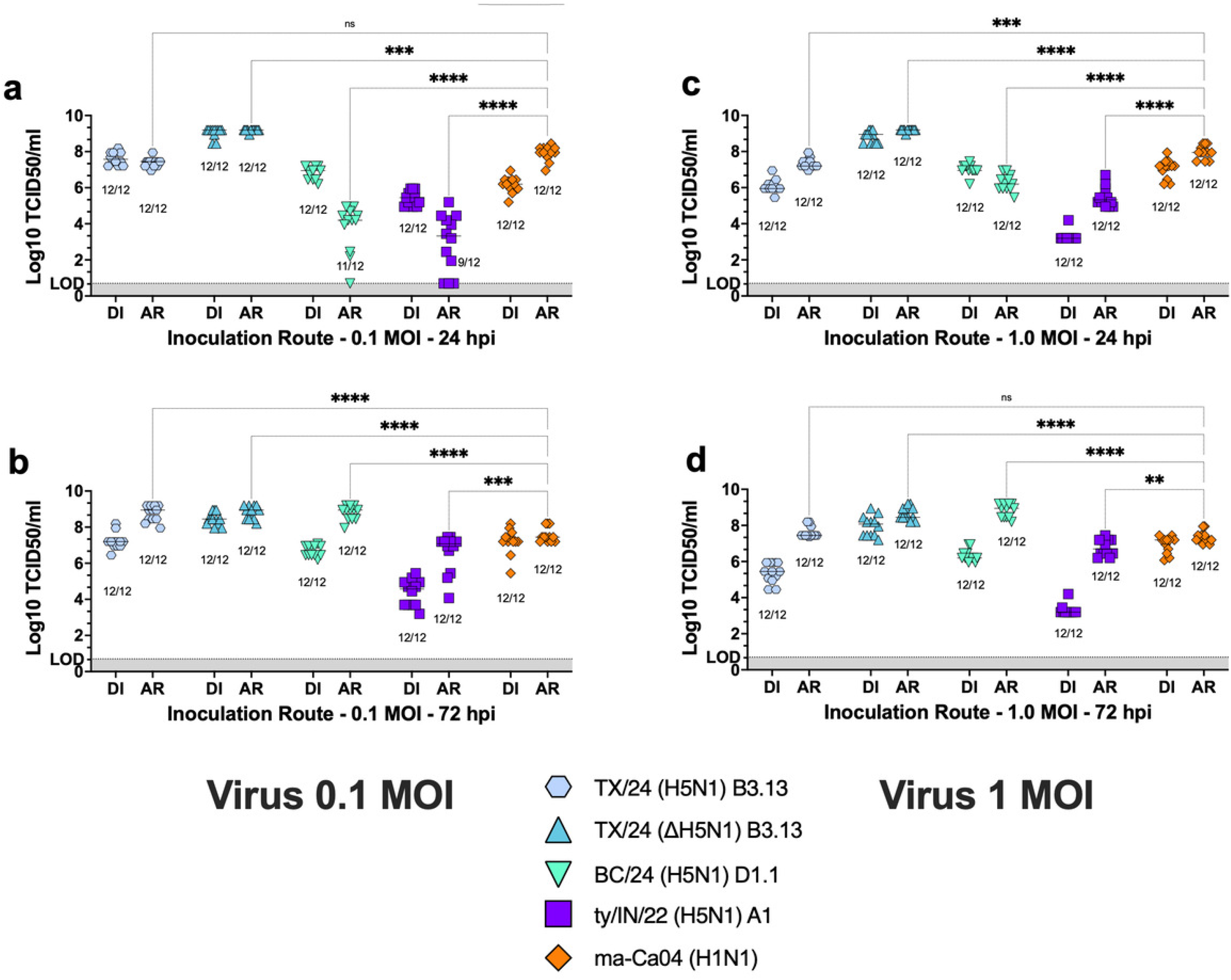
Aerosolized H5N1 viruses retain infectivity in MDCK cells. MDCK cells were infected by direct inoculation (DI) or aerosol exposure (AR) at an MOI of 0.1 **(a-b)** or 1 **(c-d)** with A/Texas/37/24 (H5N1) (TX/24), A/Texas-Halo/37/24 (H5N1) (ΔTX/24), A/British Columbia/PHL- 2032/24 (H5N1) (BC/24), A/turkey/Indiana/3707-003/22 (H5N1) (ty/IN/22), or A/California/04/2009 (H1N1) (ma-Ca04). Samples were collected at 24 and 72 h post-infection, and viral titers were determined by TCID_50_ assay. Data are presented as individual wells with median values indicated. Data were analyzed via ordinary two-way ANOVA with Dunnett’s multiple comparisons test. Full statistical details are provided in Supplementary Table 2, as some annotations were omitted from the graphs for clarity. Significance is indicated as follows: ****P < 0.0001; ***P < 0.001; **P < 0.01; *P < 0.05; ns, not significant.

### Impaired aerosol H5N1 infectivity in HAE/ALI cells does not correlate with TX/24 virulence in ferrets

Given the defective aerosol infectivity observed across all three reverse genetics-generated strains (TX/24, ΔTX/24, and BC/24), TX/24 was utilized for further characterization. We sought to determine if these in vitro defects translated to altered infectivity or pathogenesis within the ferret model (22, 26). We performed infection of both male and female ferrets (n=8/sex, n=4/group). For virus aerosol exposure of ferrets, we utilized a nose-only exposure system as previously described (37). Ferrets received 10^4^ TCID_50_/animal via aerosol or direct (liquid) inoculation routes. Animals were divided into 4 groups: aerosol-exposed females (AR-F), aerosol-exposed males (AR-M), direct inoculated females (DI-F), and direct inoculated males (DI-M). Two male ferrets in the AR-M group were humanely euthanized on day 2 post-virus exposure, although they had not reached humane endpoints, as a predetermined comparator unrelated and irrelevant to the present report. All other ferrets remained in the experimental groups and were euthanized as soon as they reached humane endpoints. Data were stratified in two ways: by route of infection (Fig. 4) and by biological sex (Fig. 5). Ferrets showed decreased activity starting at 2 dpi, with no significant variation between AR and DI groups (Fig. 4a).

**Figure 4:**
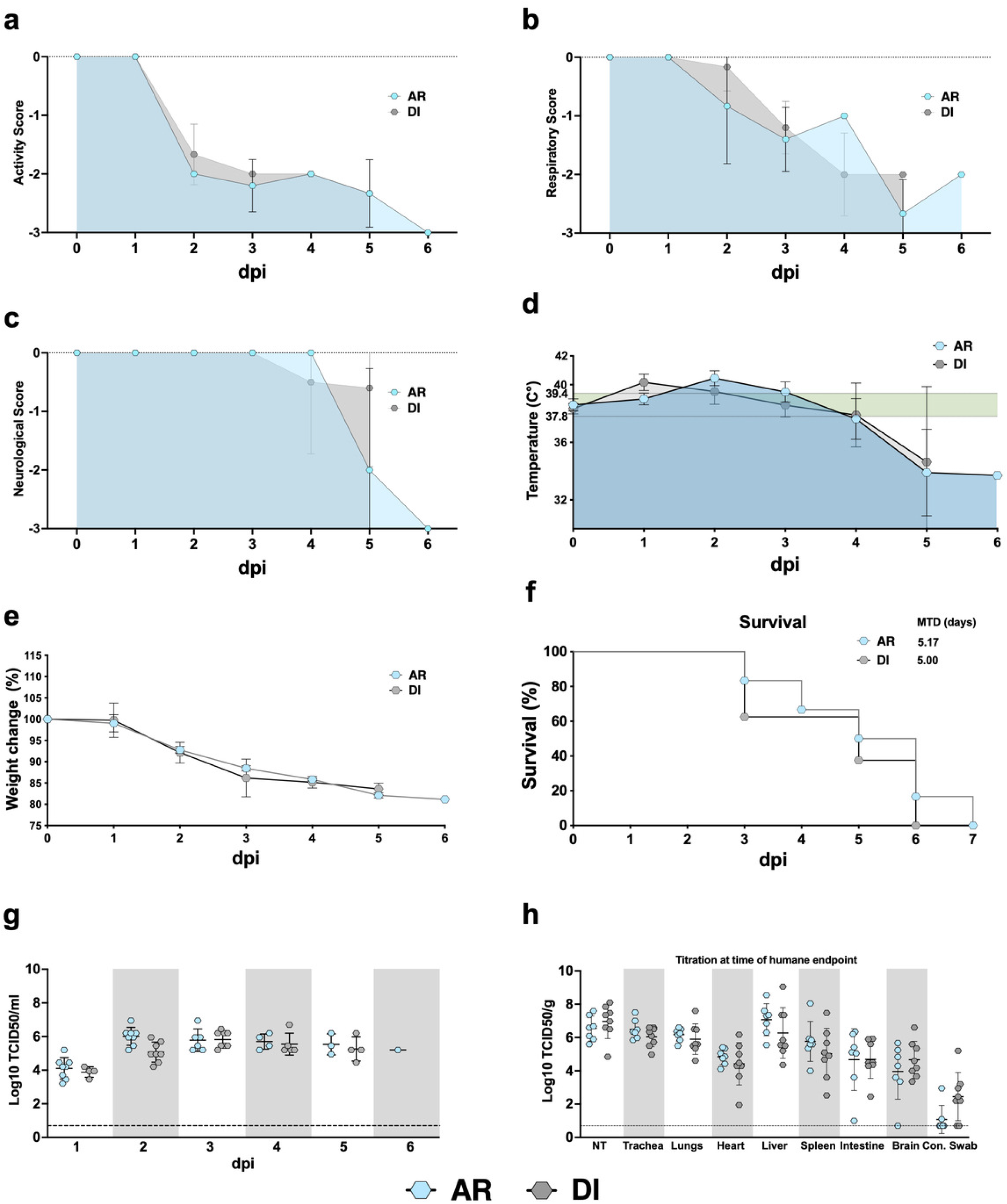
Severe disease following aerosolized and direct H5N1 infection in ferrets. Ferrets were inoculated by direct inoculation (DI) or aerosol exposure (AR) with 10^4^ TCID_50_ per ferret. Clinical signs **(a–d)**, weight loss **(e)**, and survival **(f)** were comparable between DI- and AR-inoculated ferrets. Nasal washes **(g)** were collected from 1 to 6 days post-inoculation, and viral titers were determined by TCID_50_ assay. Tissue samples **(h)** were collected at necropsy, and viral titers were determined by TCID_50_ assay. Statistical significance was assessed using multiple unpaired t-tests with BKY FDR correction **(a–e**, **g–h)** or the log-rank (Mantel-Cox) test **(f)**. Significance markers are omitted from the graphs for visual clarity; comprehensive statistical details are provided in Supplementary Table 3.

**Figure 5:**
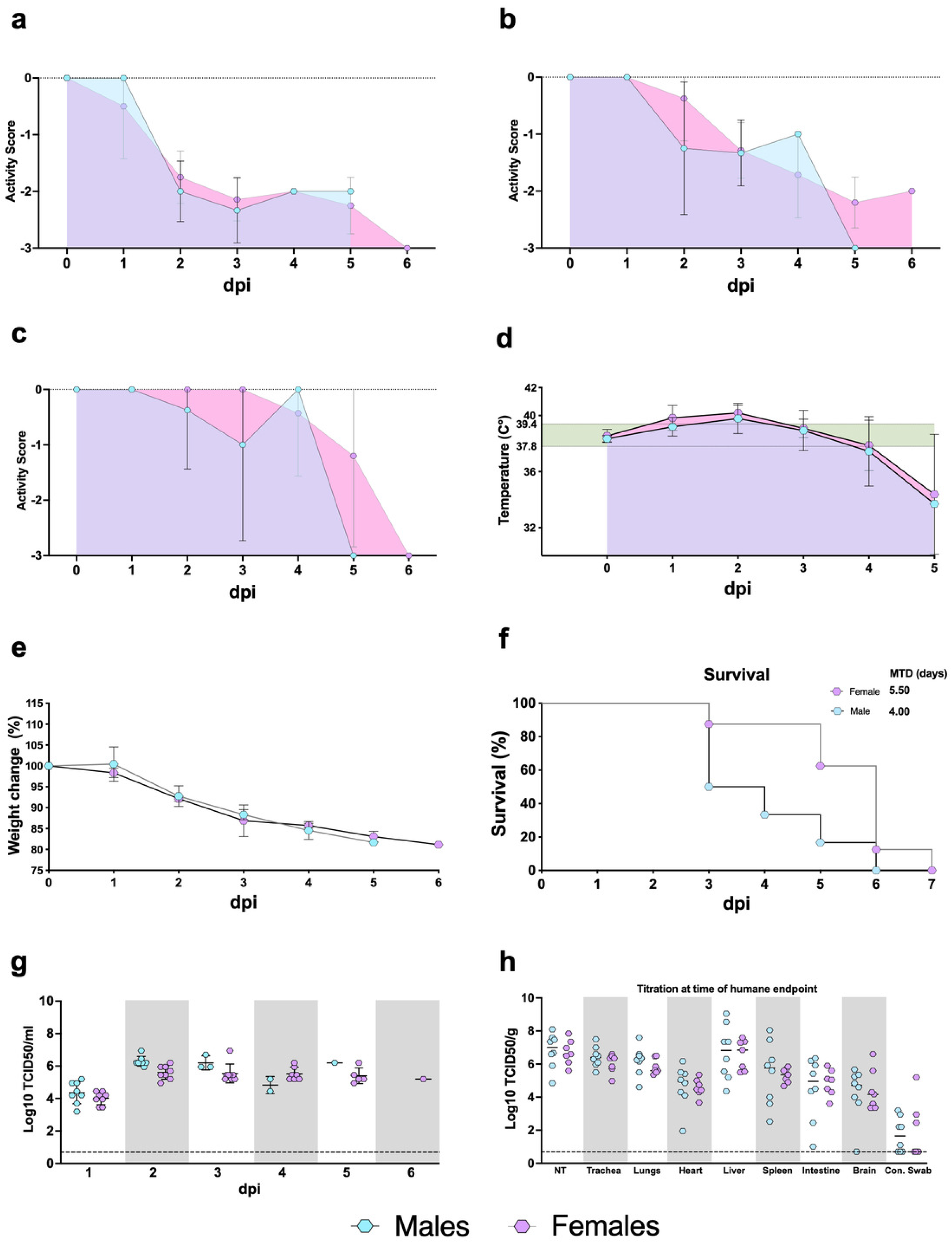
Male ferrets exhibit accelerated disease progression following aerosolized or direct H5N1 infection. Male and female ferrets were inoculated by direct inoculation (DI) or aerosol exposure (AR) with 10^4 TCID_50_ per ferret. Clinical signs **(a–d)** and weight loss **(e)** were comparable between female and male ferrets. Male ferrets succumbed to disease approximately two days earlier than female ferrets **(f)**. Nasal washes **(g)** were collected from 1 to 6 days post-inoculation, and viral titers were determined by TCID_50_ assay. Tissue samples **(h)** were collected at necropsy, and viral titers were determined by TCID_50_ assay. Data were analyzed via multiple unpaired *t*-tests with BKY FDR correction **(a–e, g–h)** and the log-rank test **(f)**. Full statistical results are in Supplementary Table 3.

Consistent with findings from a prior 2009 H1N1 study (37) comparing infection routes, TX/24- infected ferrets exhibited distinct clinical progressions based on exposure method, as AR- exposed ferrets showed more severe respiratory scores at 2 dpi than the DI-inoculated ferrets (Fig. 4b). However, while both groups experienced respiratory decline, the DI cohort demonstrated consistently higher morbidity scores throughout the infection. Neurological clinical signs became prominent only in animals surviving to day 4 or beyond (Fig. 4c). Thermal dysregulation was observed in both groups (Fig. 4d); DI ferrets reached a peak temperature increase of 1.83°C at 1 dpi, while AR ferrets peaked at 1.85°C at 2 dpi. This febrile phase was followed by hypothermia in “long-term” survivors (≥ 4 days). These clinical signs were accompanied by substantial and progressive weight loss in all ferrets with no statistically significant differences among the groups (Fig. 4e). Ultimately, both routes were lethal, with mean times to death (MTD) of 5.17 days for AR and 5 days for DI (Fig. 4f).

Analyses of nasal washes revealed robust viral shedding in both AR and DI ferrets, with peak titers of 4.9-6.9 log_10_TCID_50_/mL and 5.1-6.6 log_10_TCID_50_/mL, respectively (Fig. 4g). AR ferrets exhibited a trend toward higher shedding at 1 and 2 dpi compared to the DI group. At humane endpoints (2–6 dpi), infectious virus was detected systemically in both groups, including the nasal turbinates, trachea, lungs, heart, liver, spleen, intestines, and brain (Fig. 4h). However, while all DI ferrets (n=8) showed dissemination to the brain, 3/8 AR ferrets had undetectable viral levels in the brain. In addition, conjunctival swabs were positive in 6/8 DI and 4/8 AR ferrets.

Both male and female ferrets exhibited nearly identical physiological trajectories in response to inoculation (Fig. 5). Both cohorts demonstrated progressive lethargy, as indicated by steadily declining activity scores, and experienced rapid, uniform weight loss, reaching 80% to 85% of their initial baseline weights prior to reaching humane endpoints. Despite these similar clinical presentations, significant sex-dependent disparities were observed in overall survival (Fig. 5f). Male ferrets succumbed to the infection at an accelerated rate, exhibiting an MTD of 4 days (data calculated without considering the two male ferrets in AR-M group humanely euthanized on day 2 post-virus exposure). In contrast, female ferrets demonstrated a more prolonged survival period, maintaining an MTD of 5.5 days. Viral titers indicated robust, systemic dissemination in both sexes, with levels frequently clustering between 4 and 8 log_10_ TCID_50_/ml (Fig. 5g). Notably, the viral burdens across these tissues did not present stark discrepancies between the male and female cohorts (Fig. 5h). This suggests that the accelerated mortality observed in male ferrets may be mediated by alternative host responses or pathogenic mechanisms rather than broad discrepancies in viral replication within these evaluated tissues.

Collectively, these data demonstrate that TX/24 infection via either aerosol or direct inoculation causes high mortality and systemic spread in ferrets. These results highlight a divergence between the HAE/ALI system and the ferret model, confirming that the impaired aerosol infectivity of TX/24 previously observed *in vitro* does not reflect a generalized loss of fitness or virulence *in vivo*.

### Retention of airway mucus reduces aerosol infectivity in HAE/ALI cultures

Airway mucus serves as a frontline mucosal defense by limiting pathogen access to epithelial target cells (38). HAE/ALI cultures recapitulate this feature through the presence of mucus-producing goblet cells (33, 34). To determine whether retained airway mucus alters susceptibility to aerosol infection, HAE/ALI cultures were infected either following apical washing or with mucus retained for the 28-day ALI incubation prior to aerosol exposure at an MOI of 0.1, 1, or 10 with ma-Ca04 (H1N1) or ΔTX/24 (H5N1). MDCK cells were placed in the exposure system as described above as a control. Under mucus-retained conditions, neither ma-Ca04 nor ΔTX/24 established detectable infection at an MOI of 0.1 by 72 hpi, whereas washed cultures supported productive infection for the ma-Ca04 strain (12/12 positive wells) but less for the ΔTX/24 (H5N1) (3/12 positive wells) (Fig. 6a) concurrent with previous results under the same conditions (Fig. 2). Increasing the inoculum partially overcame this restriction, although strain-dependent differences remained evident. At an MOI of 1, productive infection was detected in only 1/12 ΔTX/24-exposed wells under mucus-retained conditions, whereas ma-Ca04 infected 12/12 wells (Fig. 6b). In contrast, all washed cultures were productively infected by both viruses. At an MOI of 10, both viruses infected all cultures irrespective of washing condition, indicating the inhibitory effect of mucus can be overcome at sufficiently high aerosol doses (Fig. 6c).

**Figure 6:**
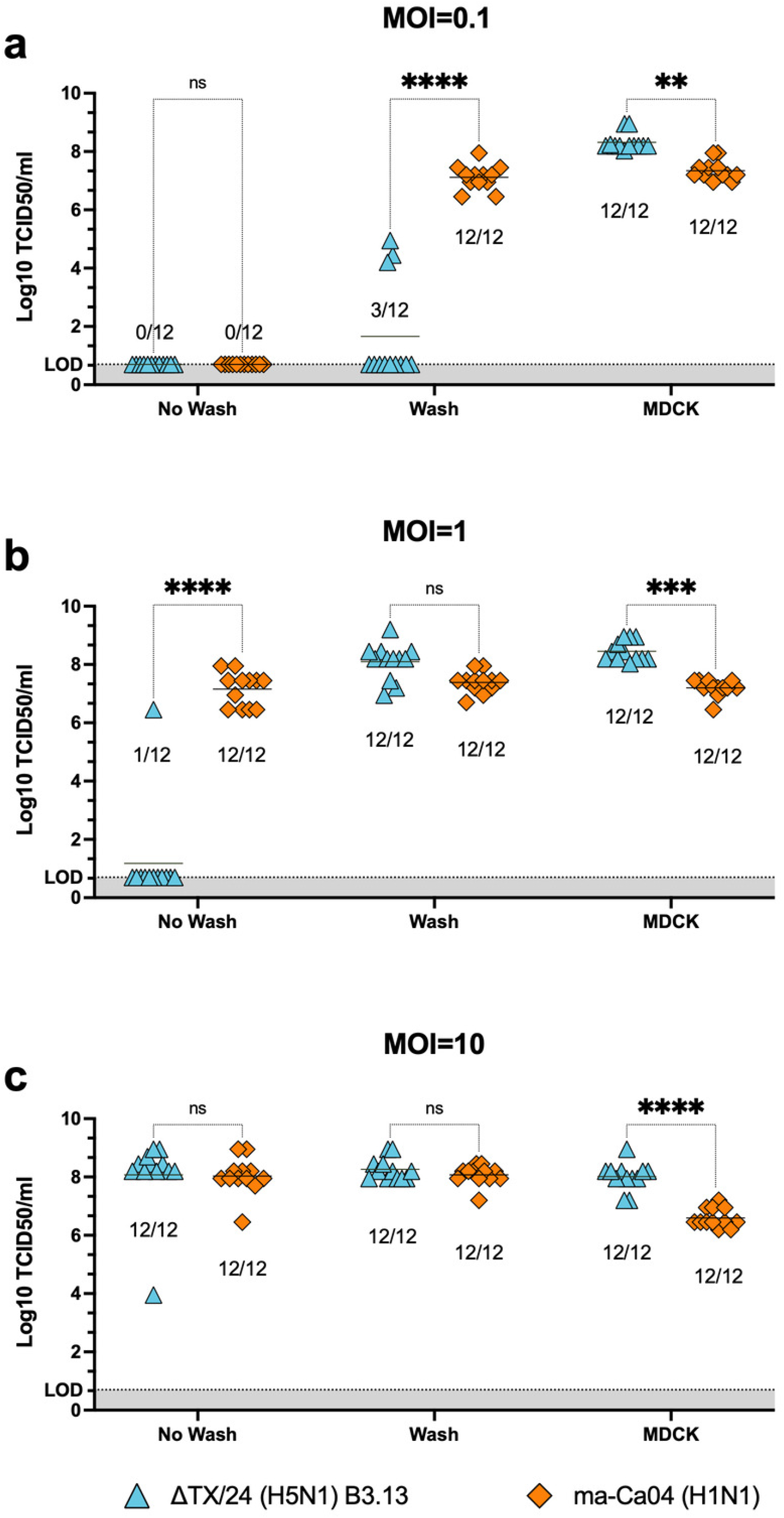
Washing HAE cells prior to aerosolized infection increases susceptibility. BCi- NS1.1 cells were either washed or left unwashed prior to infection with A/Texas-Halo/37/24 (H5N1) (ΔTX/24) or A/California/04/2009 (H1N1) (ma-Ca04) at an MOI of 0.1 **(a)**, 1 **(b)**, or 10 **(c)**. Samples were collected at 72 h post-infection, and viral titers were determined by TCID_50_ assay. Data points represent individual wells, with median values indicated. Data were analyzed using an ordinary two-way ANOVA followed by Šidák’s multiple comparisons test. Select significance indicators are displayed on the graph; full statistical results are available in Supplementary Table 2. Statistical significance depicted as follows: *P < 0.05, **P < 0.01, ***P < 0.001, ****P < 0.0001 (ns = not significant).

These findings identify airway mucus as a key barrier limiting aerosol infection in HAE/ALI cultures and suggest that efficient viral NA activity is required to overcome this restriction at low infectious doses.

### Impact of NA origin on HAE/ALI restriction

Influenza virus infectivity is contingent upon a functional balance between the surface glycoproteins, HA and NA (39–41). Throughout the ongoing panzootic, clade 2.3.4.4b H5N1 viruses have undergone continuous evolution and reassortment, resulting in high genotypic diversity (6, 7). For example, the B3.13 genotype emerged following reassortment between a B3.6 ancestor (carrying Eurasian HA and NA segments) and several internal gene segments (PB2, PA, and NP) from North American low pathogenic avian influenza (LPAI) viruses (42). Notably, while the B3.13 (and B3.2) N1 segments are of Eurasian origin, they are not direct descendants of the ancestral 1996 Goose/Guangdong (Gs/Gd) N1 lineage. Instead, they belong to a non-Gs/Gd N1 lineage acquired through reassortment with Eurasian wild bird viruses circa 2020 (12, 16).

The D1.1 genotype represents an additional distinct evolutionary shift characterized by an "NA shift." While it maintains the clade 2.3.4.4b Eurasian H5 HA, its original N1 was displaced by a North American LPAI N1 segment circulating in local wild bird populations (18). This replacement is a defining feature that distinguishes D-series genotypes from the B-series. Given these distinct avian origins, we hypothesize that the NA segments of B3.13 and D1.1, which lack extensive mammalian adaptation, contribute to the restricted ability of these viruses to infect HAE/ALI cells via the airborne route.

To determine whether differences in NA origin activities contribute to restriction of HAE/ALI infection, reassortant 1:7 H1N1 viruses containing the N1 NA of either TX/24 or BC/24 in the background of the ma-Ca04 backbone strain were evaluated under aerosol exposure at MOI 0.1 (Fig. 7a-c). At 24 hpi, both mutant viruses exhibited decreased infectivity relative to ma-Ca04, with NA:TX/24 (H1N1) infecting 3/12 wells and NA:BC/24 (H1N1) infecting 5/12 wells (Fig. 7a). By 72 hpi, H1N1:TX/24 remained restricted with 7/12 wells positive whereas H1N1:BC/24 achieved 12/12 wells positive; however, viral titers for both reassortants were significantly decreased compared to the ma-Ca04 wild-type strain (Fig. 7c). Such restriction was largely alleviated when HAE/ALI cells were exposed to an MOI of 1 of aerosolized virus (Fig. 7d-f).

**Figure 7:**
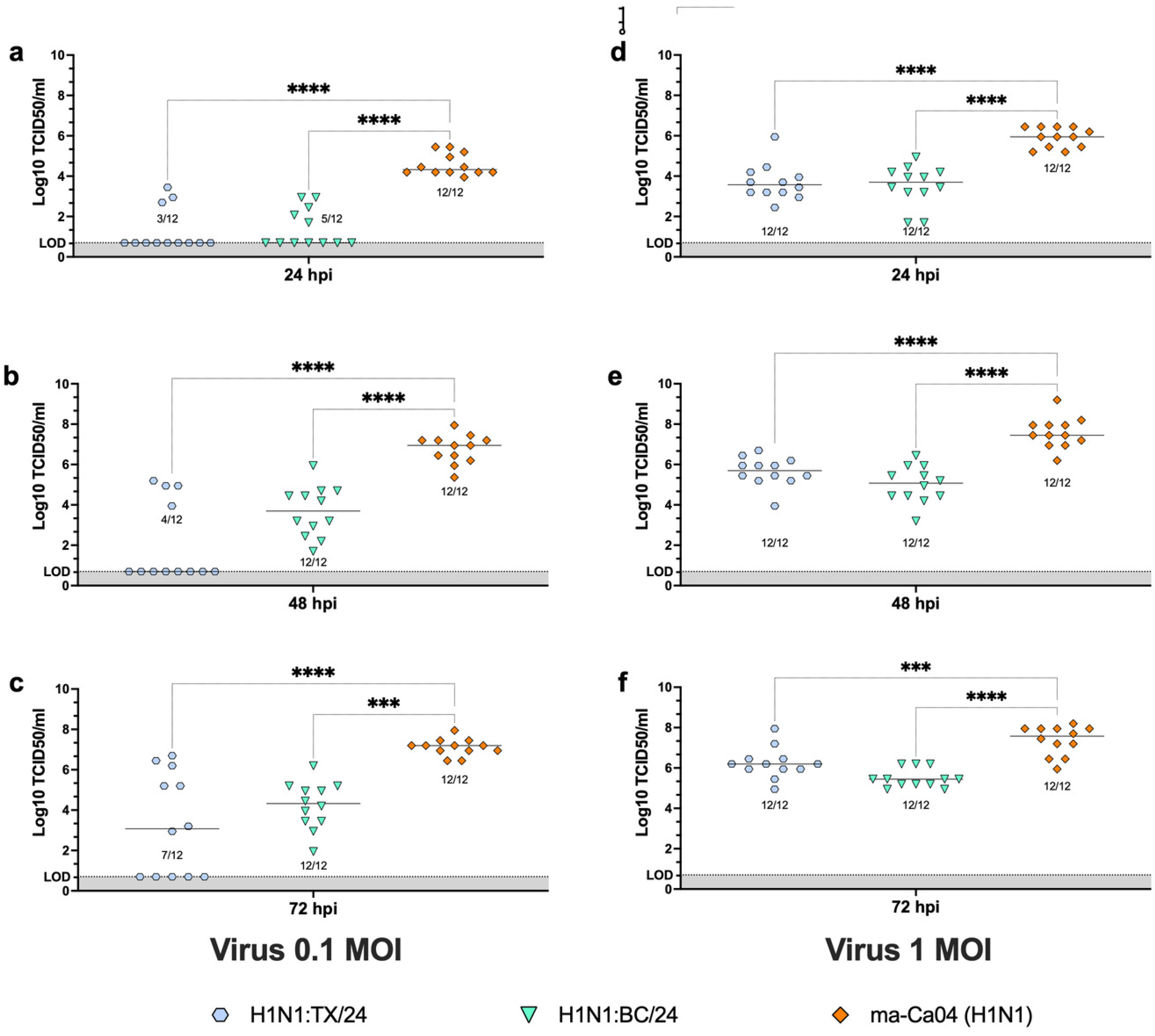
Reduced susceptibility of HAE cells to aerosolized H1N1 infection with N1 of H5N1 origin. HAE/ALI cultures were infected by aerosol exposure (AR) at an MOI of 0.1 **(a-c)** or 1 **(d-f)** with A/California/04/2009-NA Texas (H1N1) (H1N1:TX/24), A/California/04/2009-NA British Columbia (H1N1) (H1N1:BC/24), or A/California/04/2009 (H1N1) (ma-Ca04). Samples were collected at 24, 48, and 72 h post-infection, and viral titers were determined by TCID_50_ assay. Data are presented as individual wells with median values indicated. Statistical significance was assessed using ordinary one-way ANOVA followed by Tukey’s **(a, b, d–f)** or Dunnett’s T3 **(c)** multiple comparisons tests. Complete statistical analyses are detailed in Supplementary Table 2. *P < 0.05, **P < 0.01, ***P < 0.001, ****P < 0.0001; ns, not significant.

Although the kinetics of virus growth for these reassortants were slower than those of the ma- Ca04 wild-type strain in MDCK cells, both reassortants successfully initiated infections following aerosol exposure (Fig. 8a-f). These findings indicate that incorporation of the N1 NA from either TX/24 or BC/24 reduced replication efficiency of the H1N1 reassortant viruses in differentiated HAE/ALI cultures in a dose-dependent manner. Furthermore, and consistent with these phenotypes, NA activity measured using the NA-*Star* assay demonstrated a reduction in enzymatic activity (Fig. 8g). At the same concentration of virus, ma-Ca04 exhibited the highest NA activity, H1N1:TX/24 demonstrated the lowest activity, and H1N1:BC/24 showed an intermediate activity. Thus, reduced NA enzymatic activity correlated with the decreased replication efficiency in HAE cells.

**Figure 8:**
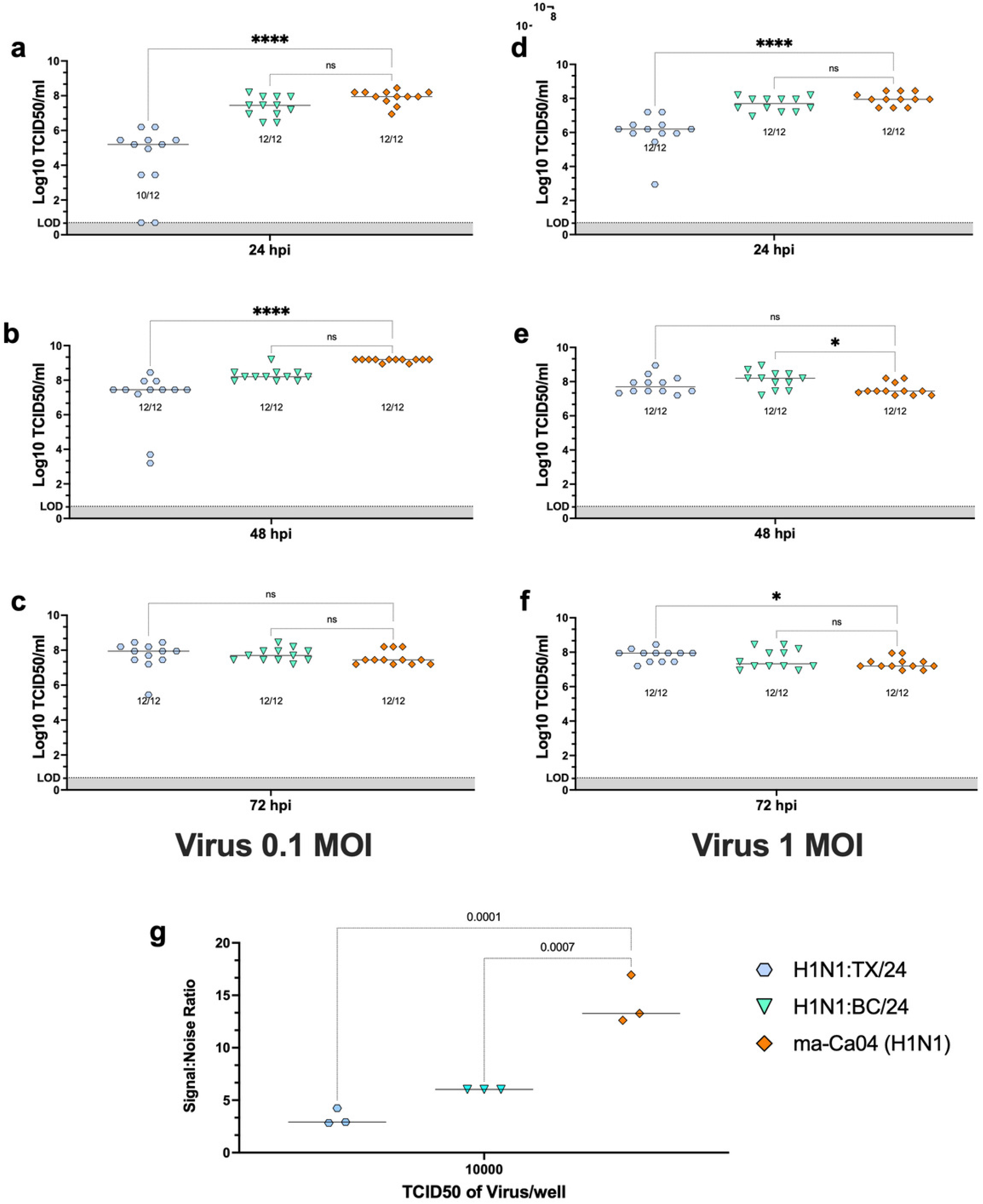
MDCK cells remain permissive to aerosolized H1N1 reassortant viruses. MDCK cells were infected by aerosol exposure (AR) at an MOI of 0.1 **(a-c)** or 1 **(d-f)** with A/California/04/2009-NA Texas (H1N1) (H1N1:TX/24), A/California/04/2009-NA British Columbia (H1N1) (H1N1:BC/24), or A/California/04/2009 (H1N1) (ma-Ca04). Samples were collected at 24, 48, and 72 h post-infection, and viral titers were determined by TCID_50_ assay. NA activity was determined using 10,000 TCID_50_ of each virus **(g)**. Data are presented as individual wells with median values indicated. Statistical significance was determined by ordinary one-way ANOVA followed by Dunnett’s T3 (**a**) or Tukey’s **(b–g)** multiple comparisons test. For visual clarity, only select comparisons are shown; comprehensive statistical analyses are detailed in Supplementary Table 2. *P < 0.05, **P < 0.01, ***P < 0.001, ****P < 0.0001; ns, not significant.

### Potential impact of polymerase complex origin on HAE/ALI restriction

Previous studies investigating the polymerase complex of clade 2.3.4.4b viruses have consistently reported reduced activity relative to contemporary human-adapted strains (43, 44). This functional disparity is attributed to specific amino acid signatures across the PB1, PB2, PA, and NP proteins (Supplementary Table 1). To evaluate polymerase function, the activity of the TX/24, BC/24, and ty/IN/22 complexes was compared at 35°C-the physiological temperature of the human lower trachea-against a contemporary 2009 pandemic H1N1 descendant, A/California/LACPHL-INF00602/2024 (CA/24). Quantitative analysis using a nanoluciferase (Nanoluc) reporter replicon, normalized to secreted alkaline phosphatase (SEAP), revealed that CA/24 consistently maintained the highest level of activity across all timepoints, peaking at 72 hours post-transfection (hpt) in HEK293T cells (Fig. 9a). While TX/24 and BC/24 exhibited intermediate, statistically distinct levels of activity (with TX/24 performing slightly higher), the ty/IN/22 complex demonstrated minimal polymerase function near baseline (Fig. 9a). These functional differences were further quantified by normalizing results against the mean CA/24 activity, which established median ratios at 72 hpt of approximately 0.475 for TX/24, 0.266 for BC/24, and 0.012 for ty/IN/22 (Fig. 9b). Independent qualitative assessment via an mCherry reporter replicon corroborated these findings; fluorescence microscopy displayed a robust, time-dependent red signal for CA/24, moderate signals for the TX/24 and BC/24 intermediates, and negligible expression for ty/IN/22, thereby confirming the hierarchical activity profile across both reporter systems (Fig. 9c). While we did not directly measure polymerase activity in HAE/ALI cells, our results in 293T cells align with prior studies (45–47). These findings suggest that polymerase activity contributes to the reduced H5N1 infectivity observed in the HAE/ALI model.

**Figure 9:**
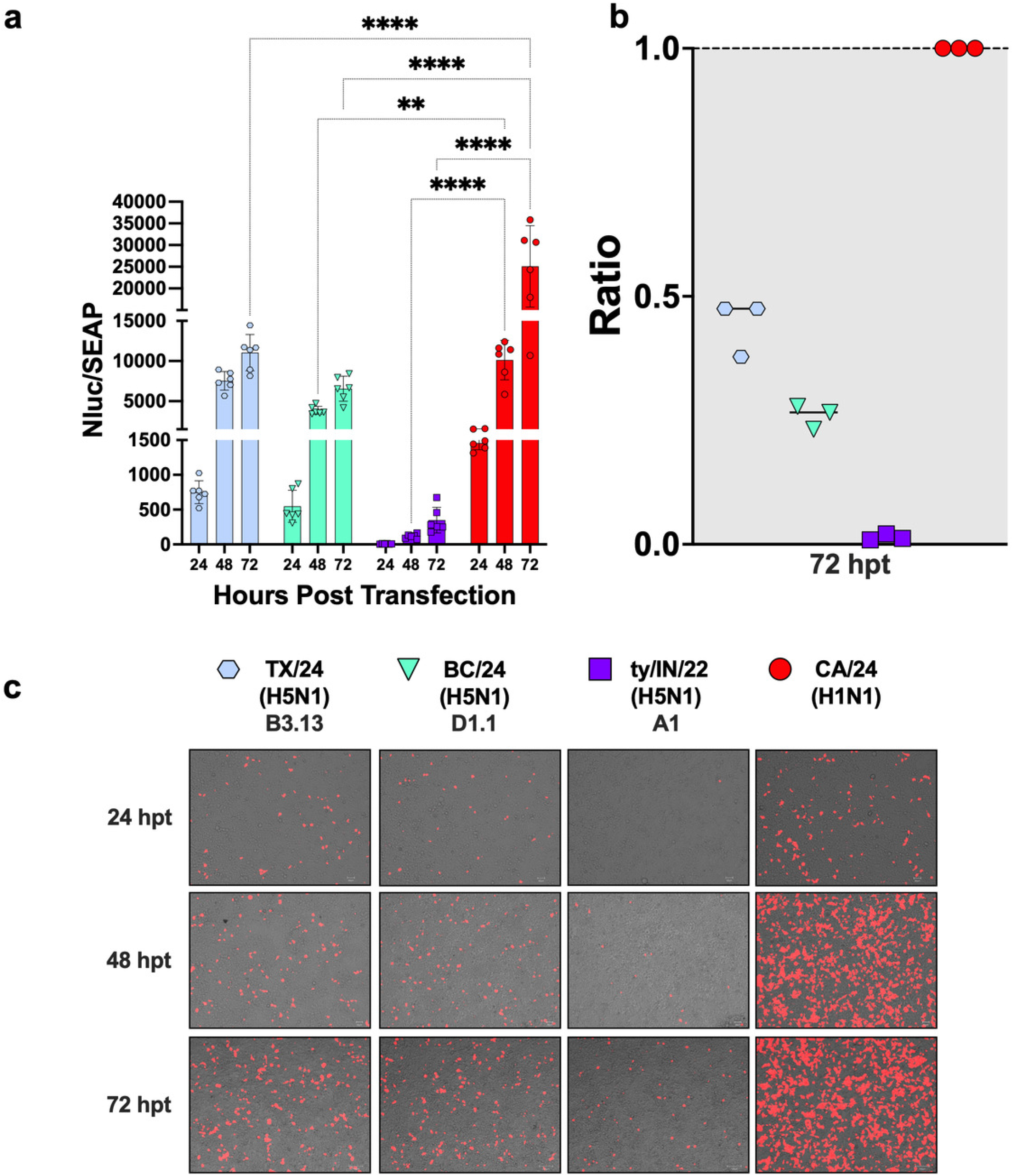
Polymerase activity of H5N1 and H1N1 complexes. Polymerase activity was assessed using a minigenome reporter assay in 293T cells. Cells were transfected with the polymerase plasmids derived from A/Texas/37/24 (H5N1) (TX/24), A/British Columbia/PHL- 2032/24 (H5N1) (BC/24), A/turkey/Indiana/3707-003/22 (H5N1) (ty/IN/22), or A/California/LACPHL-INF00602/2024 (H1N1) (CA/24), together with pIS2025_002_SecNanoLuc_A-PolI, pCMV-SEAP, and pIS2025_001_mCherry_A-PolI reporter plasmids. Samples were collected at 24, 48, and 72 hpt, and relative polymerase activity was determined by NanoLuc luminescence normalized to SEAP luminescence. Data are presented as individual wells with mean values indicated. Statistical significance was determined using ordinary 2-way ANOVA and Šidák’s multiple comparisons test (a). Relative polymerase activities are additionally shown as ratios normalized to CA/24 activity at 72 hpt (b). Representative fluorescence images demonstrating mCherry reporter expression over time are shown for each polymerase complex (c). Data were analyzed using ordinary one-way ANOVA with Tukey’s multiple comparisons test (*P < 0.05, **P < 0.01, ***P < 0.001, ****P < 0.0001; ns, not significant). Select comparisons are shown; full statistical results are available in Supplementary Table 4.

## DISCUSSION

The recent global expansion of clade 2.3.4.4b H5N1 viruses is characterized by an unprecedented host range, including highly unusual mammary gland tropism in dairy cattle and subsequent spillover into human populations (6, 7, 13). Despite this, human cases in the US have remained largely mild, predominantly presenting as conjunctivitis with limited respiratory involvement (15, 48). This clinical profile contrasts sharply with earlier Gs/Gd lineage outbreaks associated with acute respiratory distress syndrome and high mortality. Multiple investigations have examined replication of clade 2.3.4.4b viruses in differentiated HAE/ALI cultures (21, 32), yet these studies have relied predominantly on liquid inoculation. Inoculation route is a critical determinant of viral deposition, local dose distribution, and host immune activation (49). The human respiratory tract exhibits a heterogeneous distribution of α2,3- and α2,6-linked sialic acid receptors (50), and differentiated airway epithelial cultures similarly express both receptor types (33, 51). While ferret models have historically mirrored human disease progression, they are increasingly diverging from modern human outcomes. For instance, ferrets infected with contemporary seasonal H3N2 strains fail to accurately recapitulate the lower respiratory tract infections observed in humans (52, 53). Conversely, H5N1 lineages such as B3.13 and D1.1 remain notoriously lethal in ferrets, despite typically causing self-limiting illness in humans (21, 24–28, 30, 31). Our study sought to address this disconnect by comparing aerosol infectivity between these H5N1 strains and a prototypical 2009 pandemic H1N1 virus using a differentiated HAE/ALI system.

Our data demonstrate that H5N1 strains from the B3.13, D1.1, and A1 lineages exhibit a significant defect in infecting HAE/ALI cells via the aerosol route. Although this restriction is partially alleviated by increasing the viral dose or utilizing direct liquid inoculation, the HAE/ALI model consistently reflects the limited respiratory involvement observed in humans.

Interestingly, low-dose studies showed that in the few wells that became positive for H5N1— particularly the B3.13 constructs (Fig. 2a–c)—peak virus titers increased over time to levels comparable to the H1N1 control. This indicates a degree of stochasticity similar to that seen in natural human H5N1 infections. More importantly, deleting the HA polybasic cleavage site did not alter the virus’s ability to infect HAE/ALI cells, demonstrating that the site is neither beneficial nor detrimental to viral infection. Retained mucus imposed a substantial barrier to aerosol infection; however, the washed HAE/ALI condition may more closely approximate physiological airway exposure dynamics, as mucus within the conducting airway is continuously transported by mucociliary clearance and renewed on a timescale shorter than 24 hours (54). Washing the cultures 24 h prior to infection permits mucus accumulation for a limited period, which may better reflect the in vivo airway environment, where mucus is continuously cleared and replenished (54). Although the HAE/ALI cultures contain cilia for directional mucus transport, the in vitro system lacks the anatomical architecture necessary for mucus clearance comparable to that of an intact respiratory tract (55, 56). Future studies will be required to determine how mucus and its components influence infection across multiple influenza subtypes (57–59), including experiments evaluating infection following mucus removal at different time points prior to inoculation. Collectively, these findings suggest that the mucus-retained condition represents a highly restrictive environment that may exaggerate barriers to infection relative to the in vivo airway, whereas the combined use of washed and mucus-retained conditions may better bracket the range of airway environments encountered during natural exposure. In contrast, the B3.13 TX/24 strain remained highly infectious and lethal in ferrets regardless of the delivery method (aerosol vs. direct inoculation). This observation is consistent with those from several groups that demonstrated the lethality of clade 2.3.4.4b H5N1 viruses in the ferret model (21, 22); including at least one study that utilized aerosol-mediated virus delivery (26). Thus, our observations further confirm that the HAE/ALI infectivity defects do not indicate a generalized loss of fitness of the TX/24 strain but rather a model-dependent divergence. Interestingly, the D1.1 BC/24 reverse genetics strain showed highly restricted HAE/ALI infectivity. The low levels of infectivity of the D1.1 BC/24 strain were unexpected as the virus caused severe disease resulting in hospitalization (60). A likely explanation is that our reverse genetics clone lacks the mammalian-adapted mutations present in the natural isolate. However, an alternative explanation could be due to biological and/or immunological differences between the donor of the HAE/ALI cells (healthy adult male) and the hospitalized patient (preadolescent female).

Further studies beyond the scope of the present report are likely to shed more light on this apparent discrepancy.

We further demonstrate that H5N1 restriction is driven in part by the viral NA surface glycoprotein and the polymerase complex. Our reassortant H1N1 studies indicate that N1 segments derived from wild aquatic birds, which lack extensive mammalian adaptation, significantly limit HAE/ALI infectivity. Although these viruses have naturally transitioned to a respiratory tropism in birds—potentially favoring mammalian spillover—their NA enzymatic activity remains a limiting factor in human airway epithelium. Specifically, the higher NA activity observed in BC/24 reassortants compared to TX/24 may explain why some D1.1 human infections have resulted in more severe respiratory disease.

Furthermore, the polymerase complex of clade 2.3.4.4b viruses exhibited impaired activity at 35°C—the physiological temperature of the human lower trachea—relative to human-adapted strains. This finding aligns with the delayed replication kinetics observed in HAE/ALI cells.

Notably, although the HAE/ALI studies were conducted at 37°C to optimize cell growth, this temperature closer to avian physiological norms should theoretically favor H5N1 replication; however, these strains remained significantly restricted in this system.

While several polymerase mutations are established mammalian adaptation markers (61, 62)— most notably E627K and D701N in PB2—they do not fully explain the functional disparities observed in our study. Recent dairy cattle isolates have also featured a fixed M631L mutation linked to bovine adaptation (42, 46). The TX/24 strain encodes PB2 627K, and while the natural BC/24 isolate exists as a mixture of 627E and 627K, our reverse genetics clone lacks the mammalian-adapted variant. Furthermore, neither strain possesses the 701N or 631L mutations. Other signatures, such as PB2 478I and NP 450N (63), have been linked to increased virulence in ferrets, yet their role in HAE/ALI aerosol infectivity remains poorly defined.

Ultimately, these canonical markers do not account for the observed variations in polymerase activity, suggesting that additional amino acid differences within the polymerase complex contribute to these quantitative shifts (Supplementary Table 2). Finally, it is important to acknowledge that other viral factors—including the HA, M1, M2, NS1, and NEP proteins, among others—may also contribute to the restricted replication of clade 2.3.4.4b viruses within our HAE/ALI aerosol delivery system.

The marked discrepancy between high mortality in ferrets and restricted infectivity in human airway cultures suggests that traditional mammalian models may overestimate the current public health risk of clade 2.3.4.4b H5N1 viruses. While ferrets remain a gold standard for assessing systemic virulence and lethality, our findings indicate that the ferret model may not fully replicate the physiological barriers to respiratory infection encountered in the human airway. Consequently, future risk assessments should prioritize physiologically relevant platforms, such as the HAE/ALI aerosol system, to more accurately distinguish between a virus’s capacity for severe systemic disease and its potential for efficient airborne transmission in humans.

A significant advantage of this approach is the high reproducibility of the immortalized HAE/ALI system, providing a standardized baseline for initial risk characterization. This platform can be further complemented by evaluations in primary HAE/ALI cultures derived from multiple human donors to account for population-level biological diversity. Although such expanded evaluations are beyond the scope of the current manuscript, utilizing these refined *in vitro* aerosol systems will be critical for identifying the specific genetic shifts required for these viruses to overcome current barriers to human adaptation.

Beyond model selection, our data underscore that the primary obstacles to human adaptation for these viruses reside, at least in part, in the functional limitations of their NA and polymerase complexes. Consequently, future research should investigate the minimum genetic shifts required to overcome these enzymatic hurdles. Longitudinal surveillance must focus on monitoring the evolution of viral segments that could enhance replication kinetics at the human physiological temperature of 33-37°C.

Collectively, our results provide a deployable risk-assessment platform that highlights how unadapted NA and impaired polymerase activity currently limit the impact of H5N1 viruses on human health. These data also serve as a critical warning: should these circulating H5N1 strains reassort with human-adapted influenza viruses to overcome these enzymatic barriers, the risk of a new pandemic would increase substantially.

## MATERIALS AND METHODS

### Plasmids and Recombinant Viruses

The reverse genetics plasmids encoding the eight segments of the mouse-adapted H1N1 A/California/04/2009 (H1N1) (ma-Ca04) have been previously described (64). Plasmids encoding the eight wild-type and HA-Halo segments from A/Texas/37/2024 (H5N1) (65) and A/British Columbia/PHL-2032/2024 (H5N1) were synthesized by Twist Biosciences (San Francisco, CA) using publicly available sequences. Viruses were rescued with all wild-type plasmids for the A/Texas and A/British Columbia. The HA-Halo of A/Texas/37/24 was rescued with the wild-type backbone and the HA-Halo. Recombinant viruses were rescued in a 7+1 system in the background of ma-Ca04 with the NA of A/Texas or A/British Columbia. The full mouse-adapted CA04 virus served as positive control. Virus stocks were titrated by tissue culture infectious dose 50 (TCID_50_), and virus titers were established by the Reed and Muench method (66). Virus sequences were confirmed by next-generation sequencing using Plasmidsaurus sequencing services (San Francisco, CA). Viruses were generated through reverse genetics as previously described (67, 68).

### Cells

Human airway epithelial (HAE) BCi-NS1.1 cells (34) were obtained from Dr. Ronald Crystal (Weill Cornell Medicine, NY, USA). Cells were maintained in 1X Basal Media (STEMCELL Technologies, Vancouver, BC, Canada) supplemented according to the manufacturer’s instructions. Cells were cultured at 37 °C under 5% CO_2_. Differentiation of BCi- NS1.1 cells was performed on 12.5 mm Transwells with 0.4-µm-pore polyester membrane inserts (Corning Inc., Corning, NY). Before plating, Transwell membranes were coated with human type IV collagen (Sigma-Aldrich, St. Louis, MO) and then rinsed with 1X phosphate-buffered saline (PBS; Thermo Fisher Scientific, Waltham, MA). Once the membrane was dry, 300,000 BCi-NS1.1 cells/Transwell were plated with 1X Basal Media and cultured at 37 °C under 8% CO_2_. Upon reaching confluency, the cells were changed to ALI conditions by removing the apical media and then changing the basal media for 1X ALI media (STEMCELL Technologies) supplemented according to the manufacturer’s instructions. Cells were cultured in ALI conditions at 37 °C under 8% CO_2_ during early differentiation and transferred to 37 °C under 5% CO_2_ until they reached 28 days in ALI conditions.

### Virus Aerosolization

Virus aerosolization was performed in a class II biosafety cabinet under BSL3 conditions essentially as previously described with minor modifications (33). In brief, the nebulizer chamber setup contained an air pump, Buxco mass dosing controller (Data Sciences International, St. Paul, MN), Aeroneb lab control module (Kent Scientific, Torrington, CT), Aeroneb lab nebulizer unit (Small VMD; Kent Scientific), Mass dosing exposure chamber (Data Sciences International), SKC BioSampler (SKC, Eighty Four, PA), BioLite+ High-volume sample pump (SKC), SKC vacuum pump (SKC), liquid traps (Fisher Scientific, Hampton, NH), HEPA-CAP filters (VWR, Radnor, PA), and several sizes of plastic tubing and tubing adapters. The Aeroneb lab nebulizer unit was connected to the Buxco Mass dosing controller to control the nebulizer output efficiency (100%) and the duration (15 min) of aerosol generation. An input of HEPA-filtered air into the mass dosing chamber was provided by pooled air from the air pump and Buxco mass dosing controller for a combined flow rate of 16 L/min. Simultaneously, the air was sampled from the exposure chamber via the SKC BioSampler using the BioLite+ High-volume sample pump at 12.5 L/min. In a second port, virus-laden aerosols were pulled through a liquid trap and a HEPA-CAP filter by the SKC vacuum pump at 3.5 L/min. Prior to each exposure, the flow rates for each component were checked using a 4100 series flowmeter (TSI, Shoreview, MN).

Either directly before aerosolization or at the indicated time, differentiated HAE cells in a 12-well plate were washed two times with PBS before inoculation to remove accumulated mucus. Each wash consisted of pipetting the PBS five times over the cells. After that, viruses were aerosolized for HAE and MDCK cells (n=12 wells/cell type) simultaneously for 15 min with 0.5 ml of inoculum/well at the MOIs indicated. MDCK cells maintained 0.5 mL of media in the well throughout the aerosolization process so that the cells would not dry out. A subset of MDCK and HAE cells seeded and maintained under the same conditions were directly inoculated with the indicated MOIs, which were calculated based on the number of MDCK cells/well. Directly inoculated and aerosol exposed cells were incubated for 1 hr at 37 °C under 5% CO_2_.

Subsequently, the virus inoculum was removed, the cells were washed with 0.5 mL of PBS two times, and 1.5 mL of new ALI media was added to the basal compartment of a new 12-well plate and the transwells were transferred to the clean plate for the HAE cells. For MDCK cells, 1.0 mL of Opti-AB containing 1 mg/mL N-p-tosyl-ʟ-phenylalanine chloromethyl ketone (TPCK)- treated trypsin (Worthington Biochemicals, Lakewood, NJ) was added to each well.

For sample collection at 0-, 24-, 48-, and 72 hours post-inoculation (hpi) from HAE cells, 200 µL of Opti-AB was added to the apical portion of the well, and then cells were incubated for 10 min at 37 °C under 5% CO_2_. The media was then collected and stored at -80°C. For MDCK, 200 µL of tissue culture supernatant was collected (replaced with an equal volume of Opti-AB containing 1 mg/mL TPCK-treated trypsin) and stored at -80°C. All samples were titrated by TCID_50_ following the Reed and Muench method (66). All of these exposures were performed under BSL3 conditions except for the ma-Ca04.

### Ferret experiments

Ferret studies were approved and conducted in compliance with all the regulations stated by the Institutional Animal Care and Use Committee (IACUC) of the University of Georgia (AUP 2022 04-026-A3). Studies were conducted under ABSL-3 conditions at the Animal Health and Research Center. Animal studies and procedures were performed according to the Institutional Animal Care and Use Committee Guidebook of the Office of Laboratory Animal Welfare and PHS Policy on Humane Care and Use of Laboratory Animals. Animal studies were carried out in compliance with the ARRIVE guidelines (https://arriveguidelines.org). Twenty-week-old male and female ferrets were acquired from Triple F Farms (Gillett, PA). One day after arrival, ferrets were anesthetized with an i.m. injection (0.5 ml/kg) of a ketamine cocktail (20 mg/kg Ketamine, 1 mg/kg Xylazine) in the thighs, followed by blood collection to test serum samples for prior IAV infection using NP ELISA (IDEXX, Westbrook, ME). All ferrets tested negative for previous IAV exposure. A subcutaneous implantable temperature transponder (BMDS, Seaford, DE) was inserted in the back of the neck to ID and monitor the body temperature of each ferret. Ferrets were acclimated for seven days upon arrival before virus inoculation.

Ferrets were inoculated through aerosolized virus exposure using the *in vivo* nose-only aerosol exposure system that included a stackable inhalation tower with seven ports (Data Sciences International), nose-only ferret restraints with ally neck restraints (Data Sciences International), and complemented with the Buxco mass dosing controller, the Aeroneb lab control module, the Aeroneb lab nebulizer unit, HEPA-CAP filters, and several sizes of plastic tubing and tubing adapters, as described above. Five ports on the stackable inhalation tower were closed, while two ports were connected to the nose-only ferret restraints. Nebulized virus exposure was conducted at room temperature (20-22 °C) and 50-60% relative humidity. Ferrets were anesthetized as previously described before exposure. During each inoculation, two ferrets of the same sex were exposed concurrently to 1 x 10^4^ TCID_50_ of A/Texas/37/2024. In all cases, virus was nebulized for 10 min. Throughout the exposure, 5 L/min of HEPA-filtered air was inputted by the Buxco mass dosing controller to maintain an input of air that was more than doubled the resting respiratory minute volume of both ferrets. The flow rate of the inputted air was measured at the start of each experimental exposure. For each exposure, 5 mL of virus diluted in 1X PBS was aerosolized using the Aeroneb lab nebulizer with an expected particle size of 2.5-4 µm at a rate of 0.409 mL/min. Synchronously, exhaled air was removed through two outflow tubes connected to a HEPA filter. Virus inoculum was calculated for males and females by multiplying the desired final exposure concentration (1 x 10^4 TCID_50_; C_final_) by the input of air into the system (5 L/min; Q_air_) and then dividing the product by the respiratory minute volume (MV) that was estimated using the following calculation:

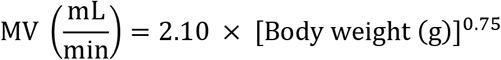

multiplied by the total exposure time (10 min; t) and the flow rate of the nebulizer (0.409 mL/min; Q_neb_).

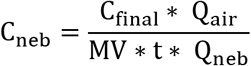

After aerosol exposure, the nose of each ferret was cleaned to limit inoculation and transmission of alternate routes. A subset of ferrets in each experiment was liquid-inoculated with the same concentrations described above in 1 ml PBS. Inoculum was instilled drop by drop into the ferret’s noses after anesthesia. Clinical signs, temperature, and weight loss were monitored daily. Nasal washes were collected daily between 1- and 7 dpi by anesthetizing the ferrets with isoflurane and administering 1 mL of PBS-BSA 1% into the nose to induce sneezing. Expelled fluid was collected into a petri dish and collected with the addition of 1 mL of PBS-AB. Once ferrets reached the humane endpoint, animals were humanely euthanized by intravenous injection of 1 mL of Euthasol (Virbac, Westlake, TX) and nasal turbinates, trachea, lungs, heart, liver, spleen, intestine, brain, and conjunctiva swab were collected.

### Tissue homogenate preparation

Tissue homogenates collected from ferrets were generated using the Tissue Lyzer II (Qiagen, Germantown, MD). Briefly, 1 mL of PBS-AB was added to each tissue with 3-mm tungsten carbide beads (Qiagen). Samples were homogenized for 15 min and then centrifuged at 15,000 g for 10 min at 4°C. Supernatants were collected, aliquoted, and stored at -80°C until further analysis. Samples were titrated by TCID_50_, and virus titers were established by the Reed and Muench method (66).

### NA Activity

NA-*Star*™ Influenza Neuraminidase Inhibitor Resistance Detection Kit (Thermo Fisher Scientific) was used following manufacturer’s instructions. Briefly, the recombinant ma- Ca04 NA viruses and the ma-Ca04 were diluted to 10,000 TCID_50_ per 50µL in NA-*Star* Assay Buffer. Triplicates of virus were added to NA-*Star* Detection Microplates and incubated at 37 °C for 10 minutes. NA-*Star* substrate was added to each well and the plate was incubated at room temperature for 10 minutes. NA-*Star* accelerator was added to each row directly prior to reading the luminescence in a plate reader. Background control wells were averaged and subtracted prior to analysis.

### TEER Measurements

Transepithelial electrical resistance (TEER) measurements were performed to assess epithelial barrier integrity of differentiated HAE/ALI cultures. Prior to measurement, 500 µL of PBS was added to both the apical and basal compartments of each Transwell insert. Resistance was measured using an EVOM2 voltohmmeter (World Precision Instruments, Sarasota, FL) according to the manufacturer’s instructions. Blank resistance values were obtained from cell-free coated Transwell inserts containing PBS alone and subtracted from sample measurements. Final TEER values (Ω·cm²) were calculated by multiplying the corrected resistance values by the membrane surface area (1.12 cm²).

### Minireplicon assay

HEK293T cells were seeded at 300,000 cells/well in a 24-well plate in DMEM (Sigma-Aldrich) supplemented with 10% FBS (Sigma-Aldrich), 1% L-glutamine (Sigma-Aldrich), and 1% penicillin/streptomycin/amphotericin B (Sigma-Aldrich) the day prior to transfection. Cells were transfected with 100 ng of each polymerase plasmid from either A/Texas (H5N1), A/British Columbia (H5N1), A/ty/IN/22 (H5N1), or A/California/LACPHL- INF00602/2024 (H1N1) (PB2, PB1, PA, and NP), and 100 ng of each of the following plasmids: pIS2025_002_SecNanoLuc_A-Pol I, pIS2025_001_mCherry_A-Pol I, and pCMV-SEAP. TransIT-LTI (Mirus, Madison, WI) was used as a ratio 2:1 (2 µL of TransIT per 1 µg of DNA) for the transfection agent. After a 45 min incubation, 100 µL of transfection reaction was added to each well (2 wells/polymerase complex). Supernatants were collected and pictures were taken 24-, 48-, and 72- h post transfection to assess the levels of NanoLuc, SEAP, and mCherry. Nluc was measured using the Nano-Glo® Luciferase Assay System (Promega, Madison, WI) and following manufacturer’s instructions. SEAP was measured only at 24-hpt using Invitrogen™ Phospha-Light™ SEAP Reporter Gene Assay System (Invitrogen, Thermo Fisher Scientific) and following the manufacturer’s instructions. Relative polymerase activity was calculated by dividing the NanoLuc activity by the SEAP activity at 24-hpt.

### Statistical analysis

All statistical analyses were performed using GraphPad Prism v10 (Boston, MA). To maintain visual clarity in the figures, the statistical significance is displayed for a subset of values. Additional statistical comparisons have been omitted from the primary figures and are provided in Supplementary Tables 2-4.

## ACKNOWLEDGEMENTS

We thank Jazmin Destiny Lynn, Hannah Walker, Karly Pecua, Morgan George, and Robert Gafnea at the Animal Health and Research Center, University of Georgia, for their assistance during animal studies under Biosafety Level 3 containment. We would also like to thank Klaudia Chrzastek and Lei He for assistance with virus titrations under Biosafety Level 2 containment.

## AUTHOR CONTRIBUTIONS

DRP developed the original aerosol exposure design. DRP and LCG designed the experiments. LCG, DR, FCF, CJC, and TSM conducted the in vivo experiment and analyzed the data. LCG, DR, FCF, IS, AL, and CJC conducted the in vitro experiments and analyzed the data. LCG and DRP interpreted the results, analyzed the data and wrote the manuscript. MS, RM, AGS, DRP edited the manuscript. All authors read, provided intellectual input and approved the final version of the manuscript.

---

**SUPPLEMENTARY TABLE 1.**
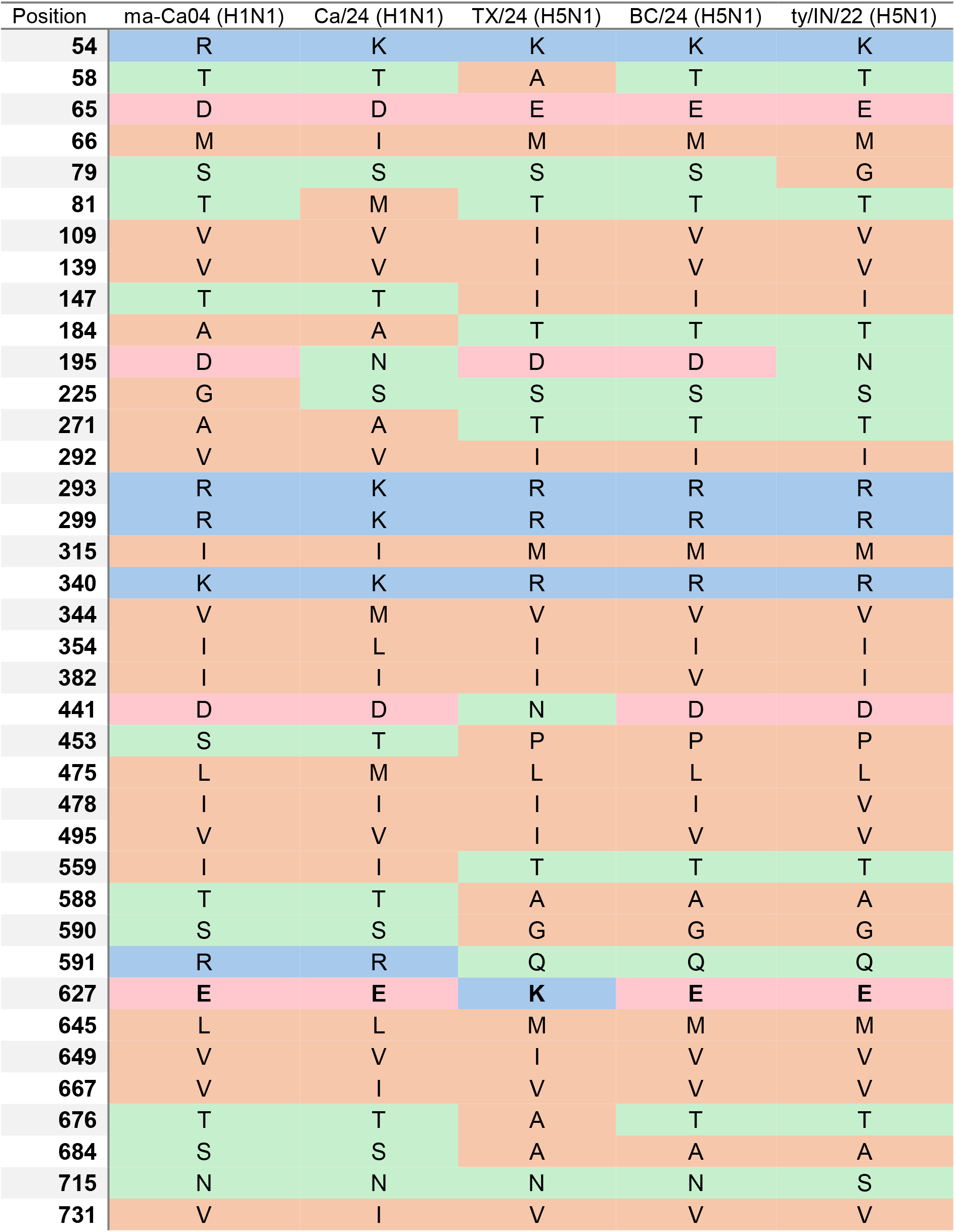
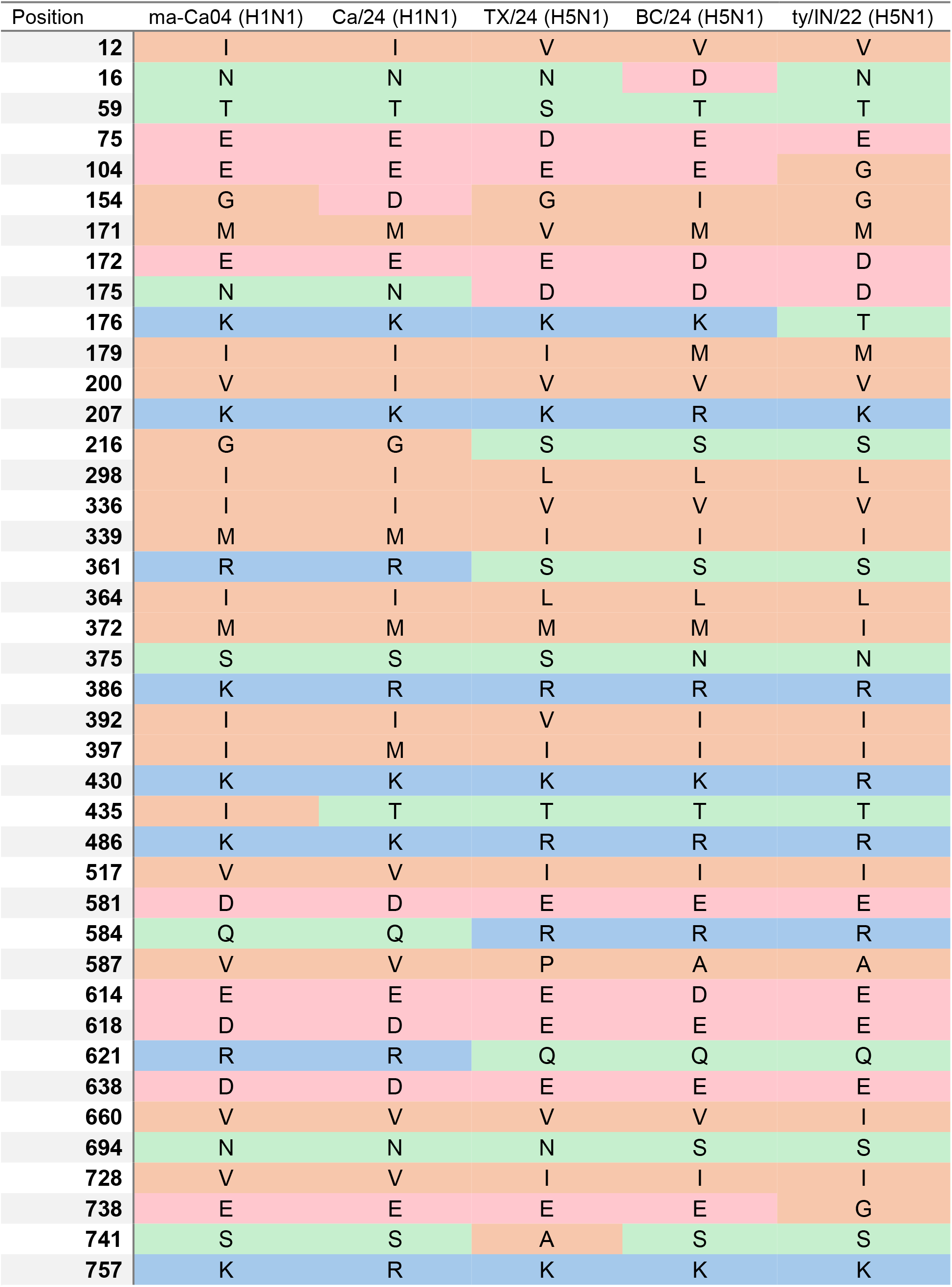
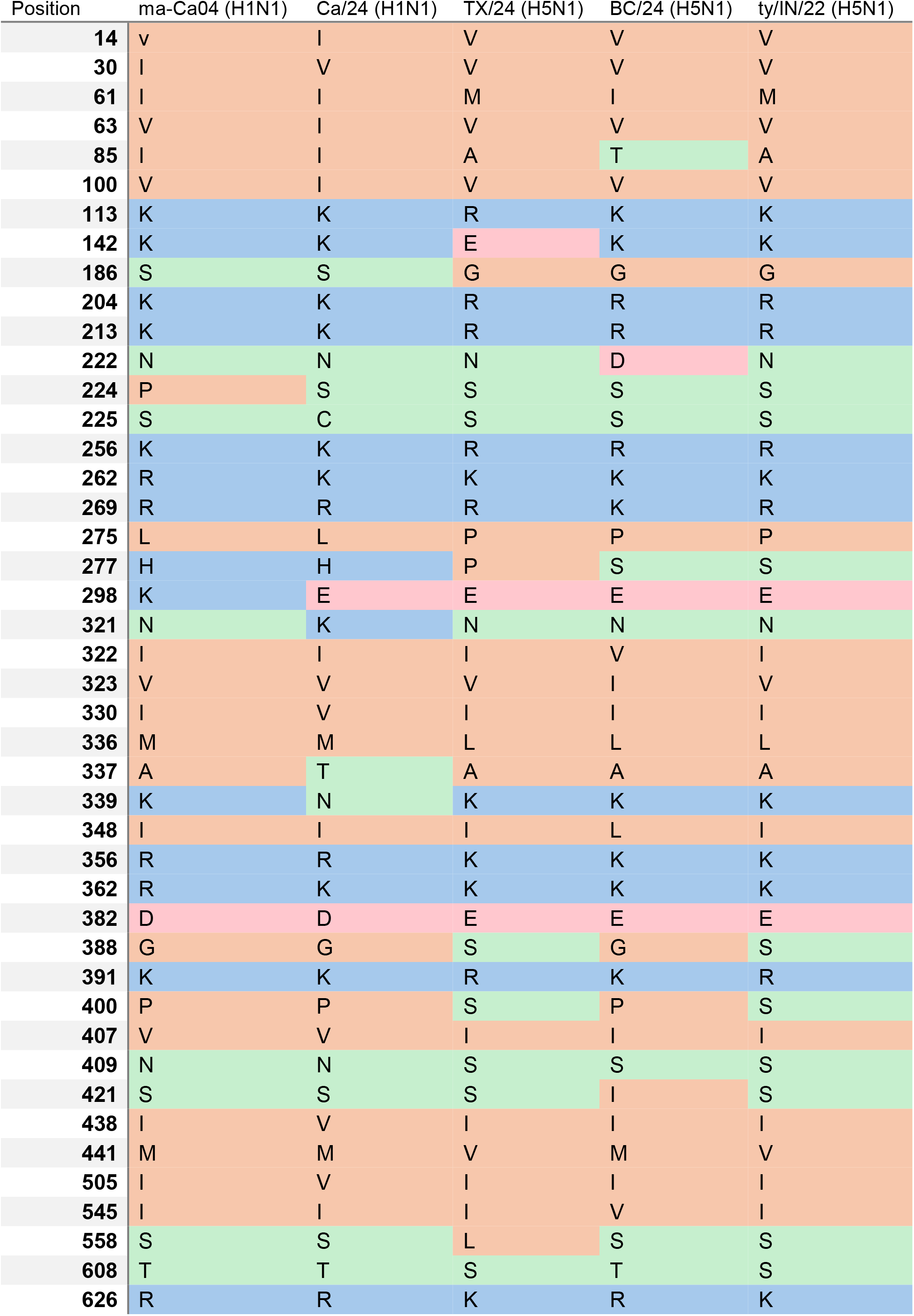
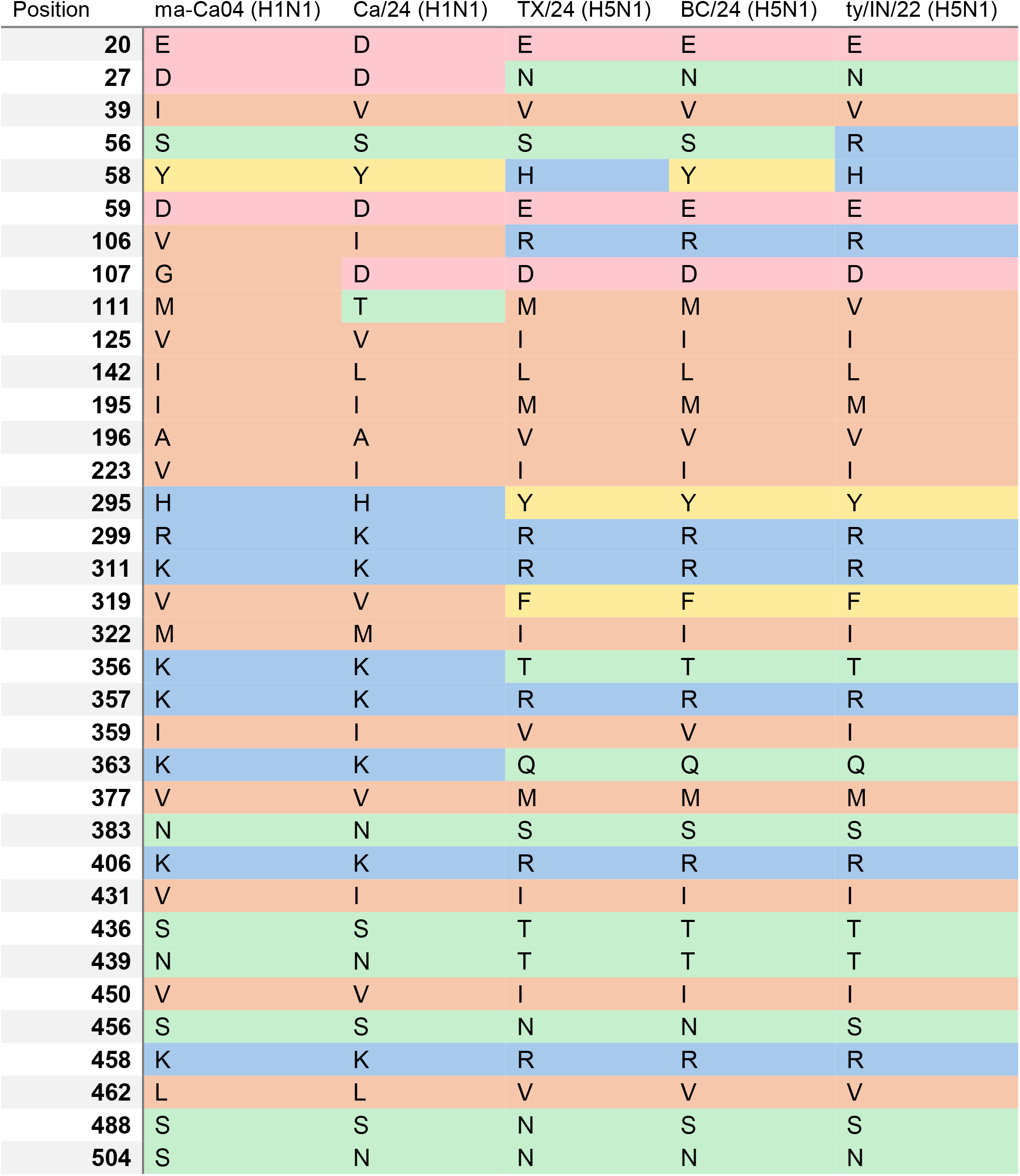
Amino Acid Variations in the Polymerase Complex (PB2, PB1, PA) and Nucleoprotein (NP) Among Selected H1N1 and H5N1 Virus Strains.

**SUPPLEMENTARY TABLE 2.**
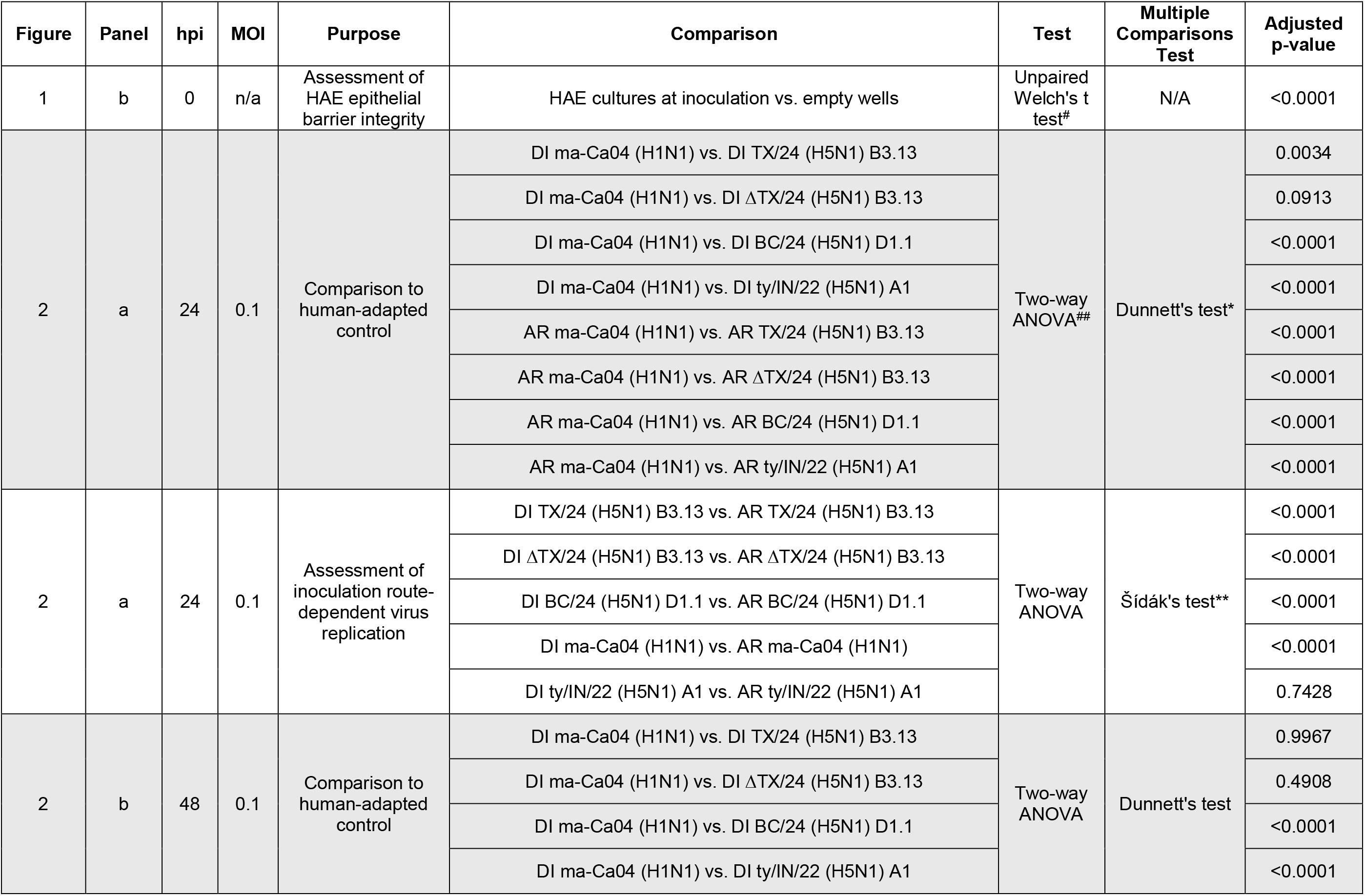
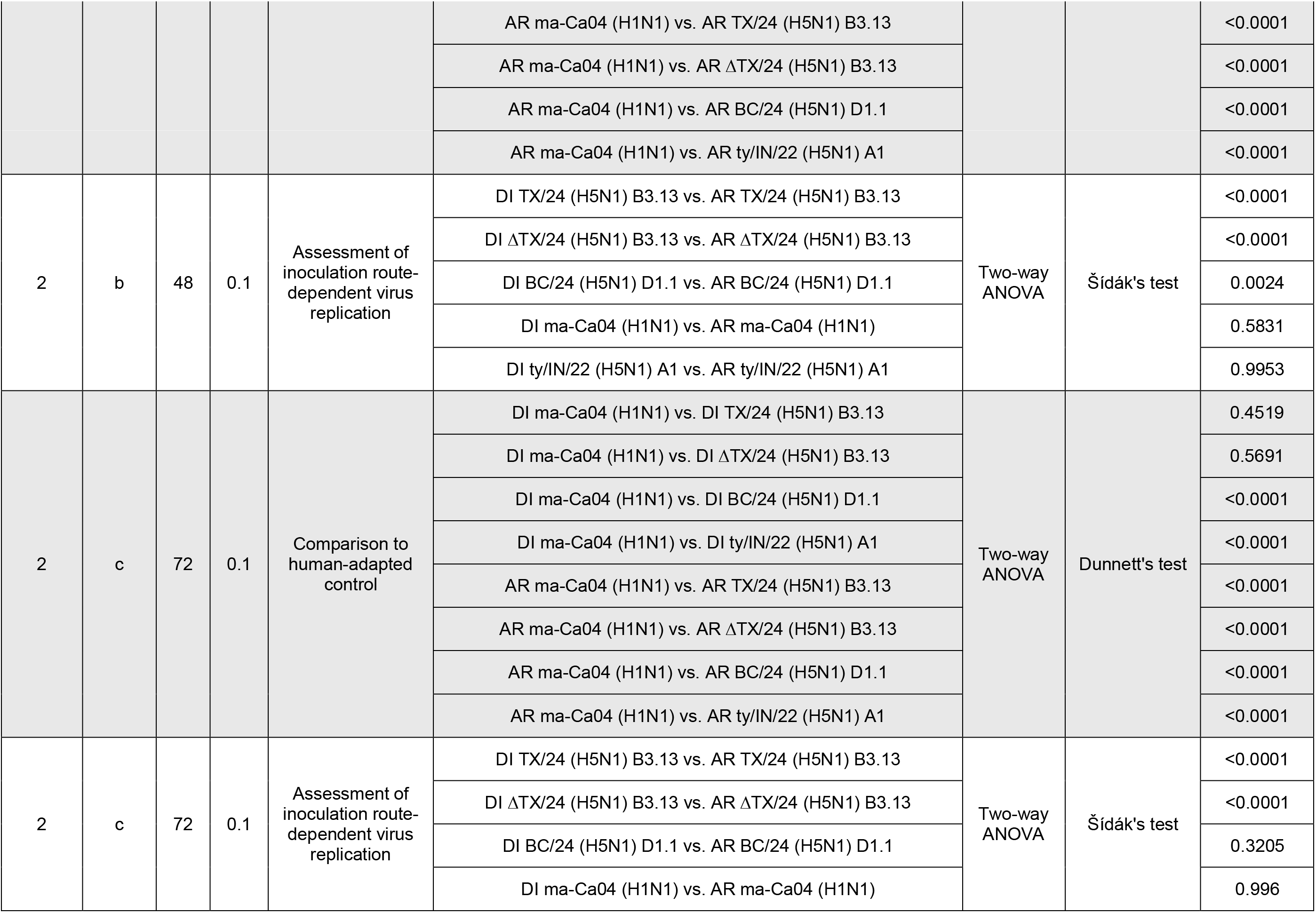
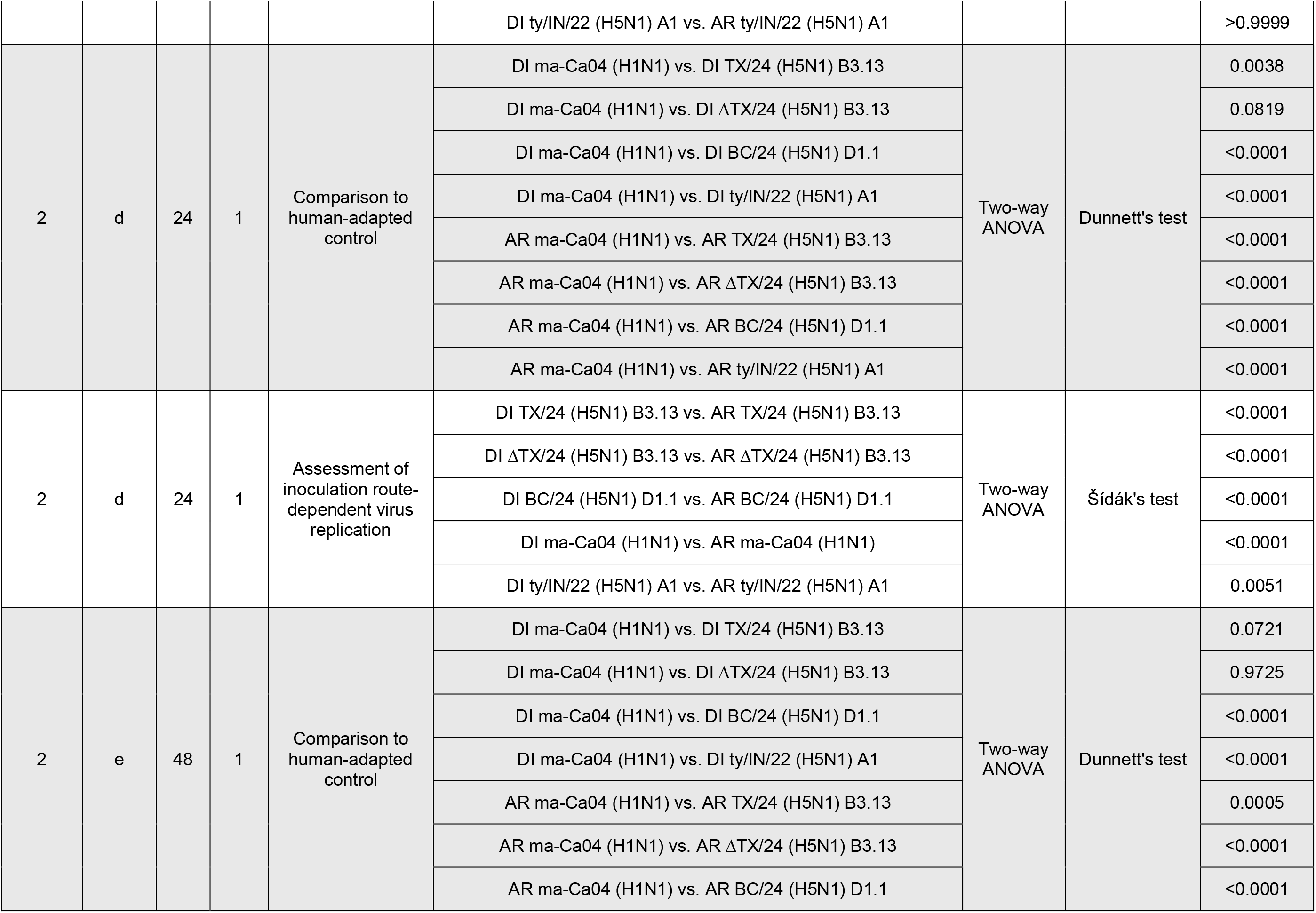
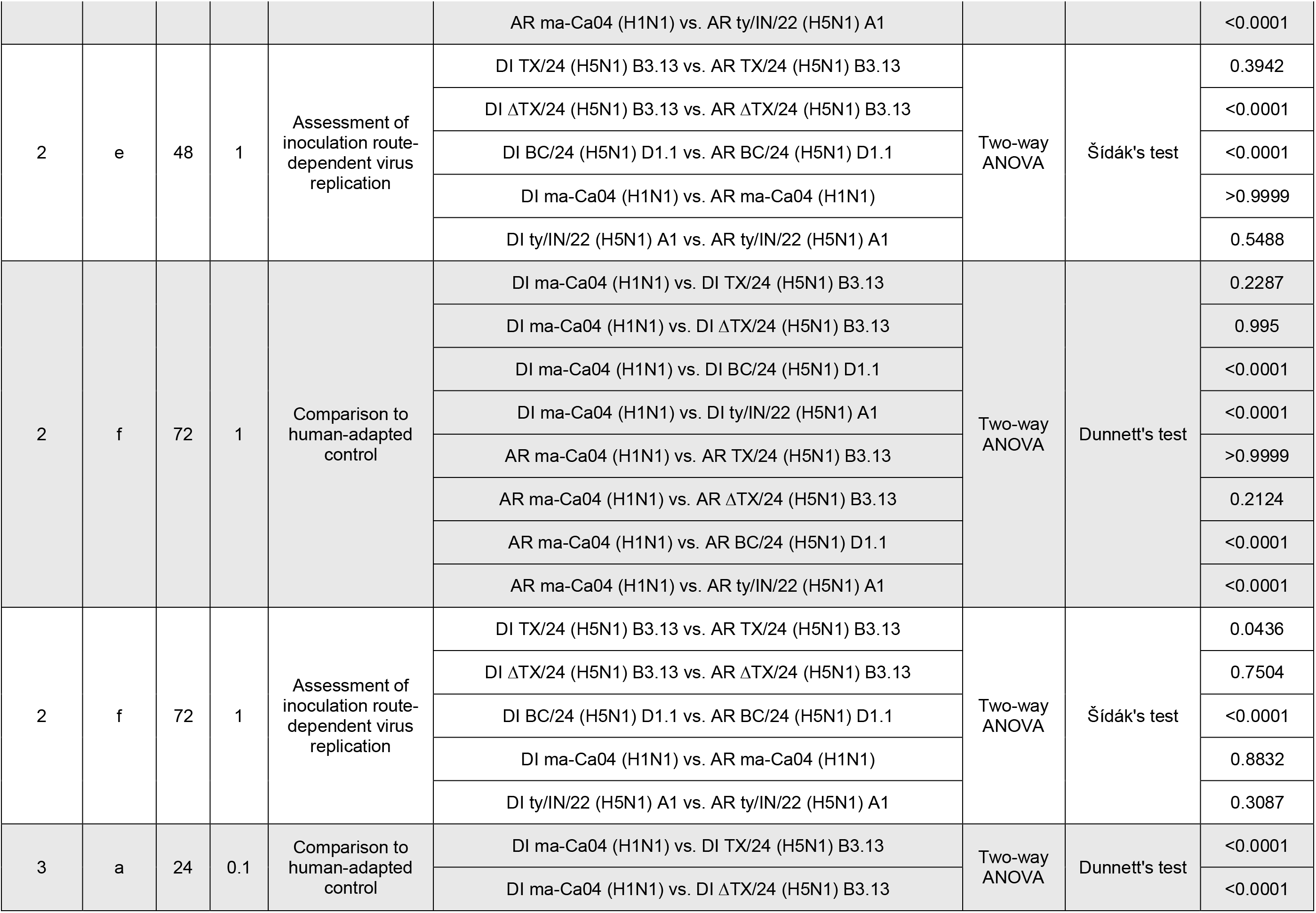
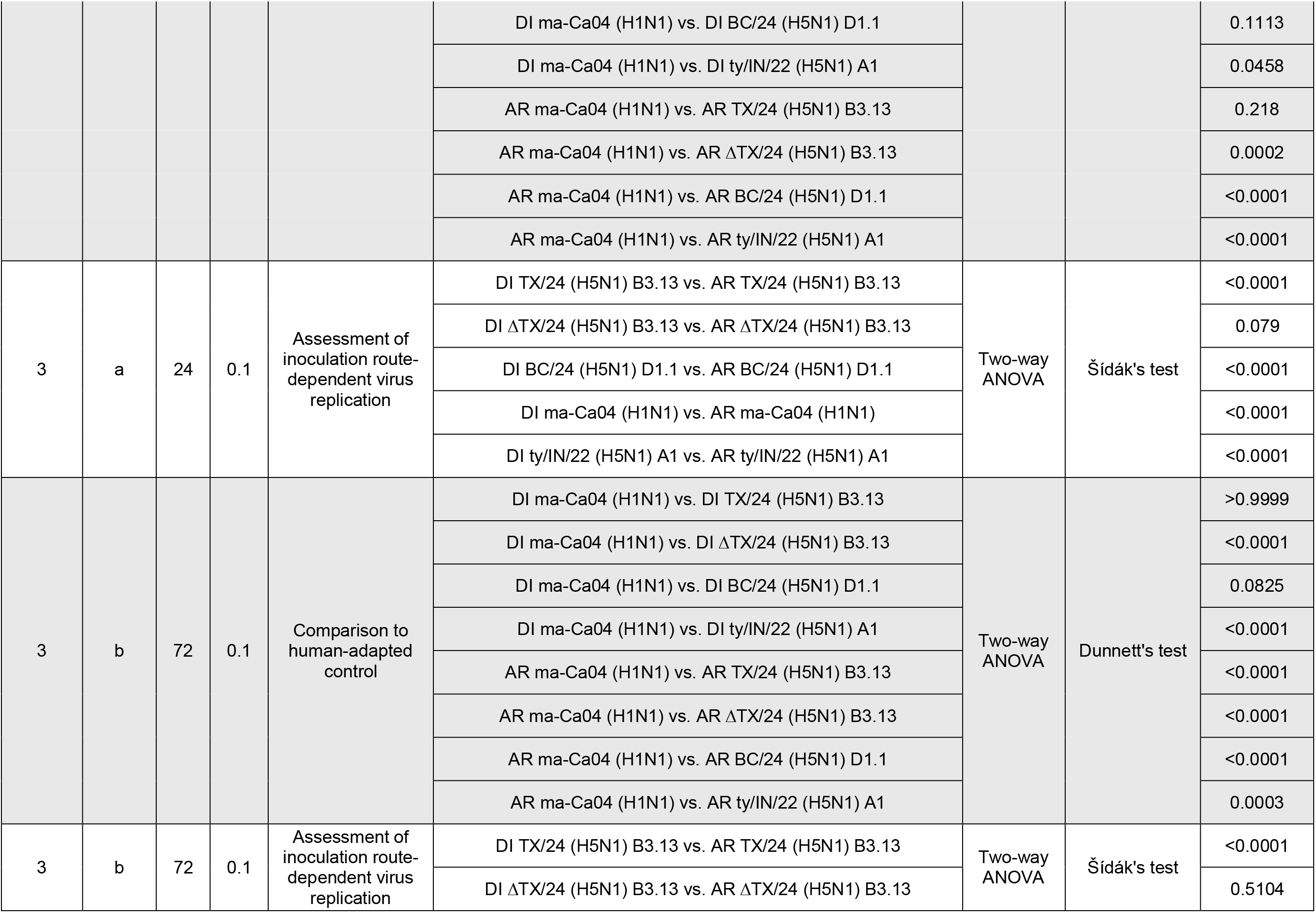
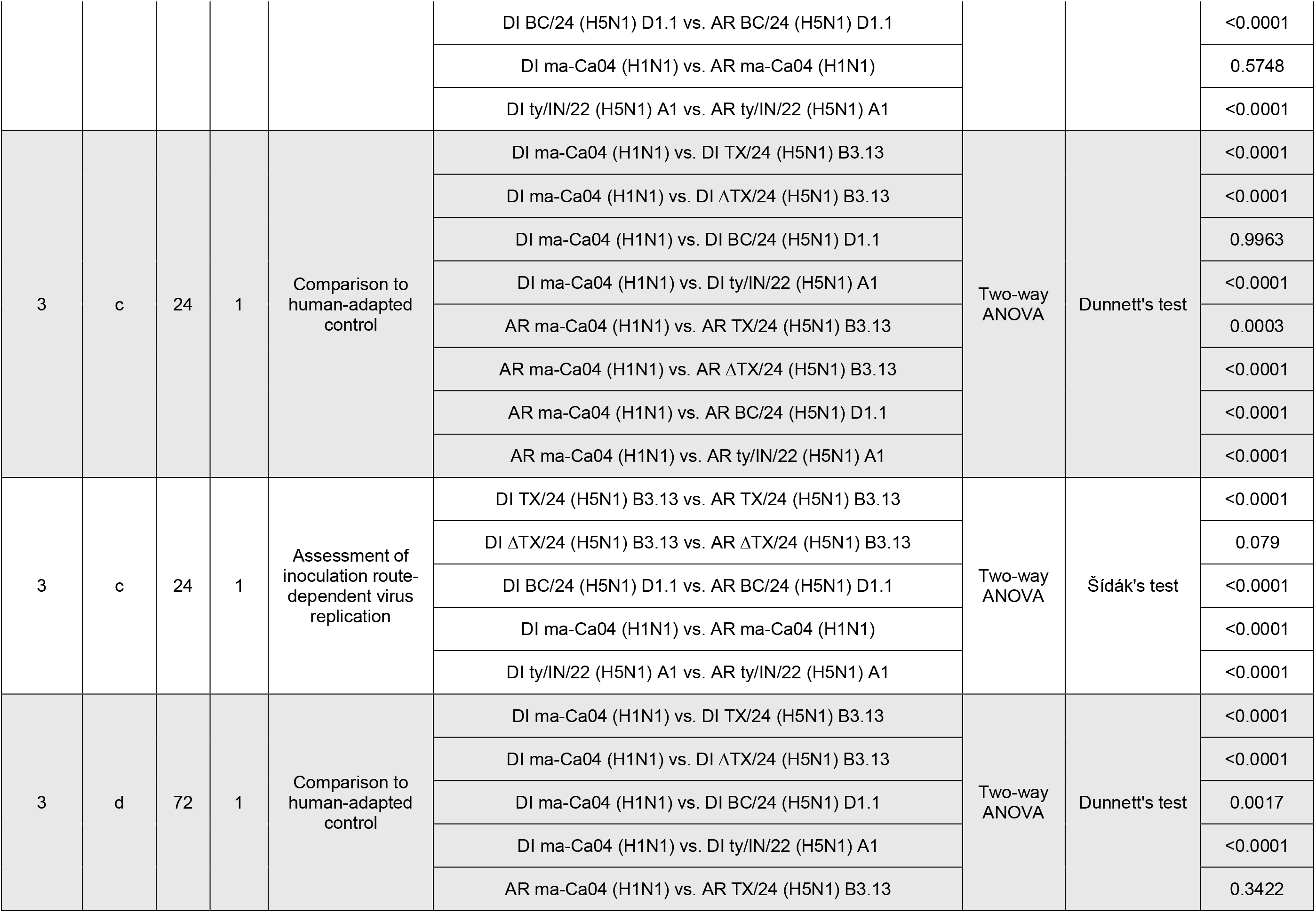
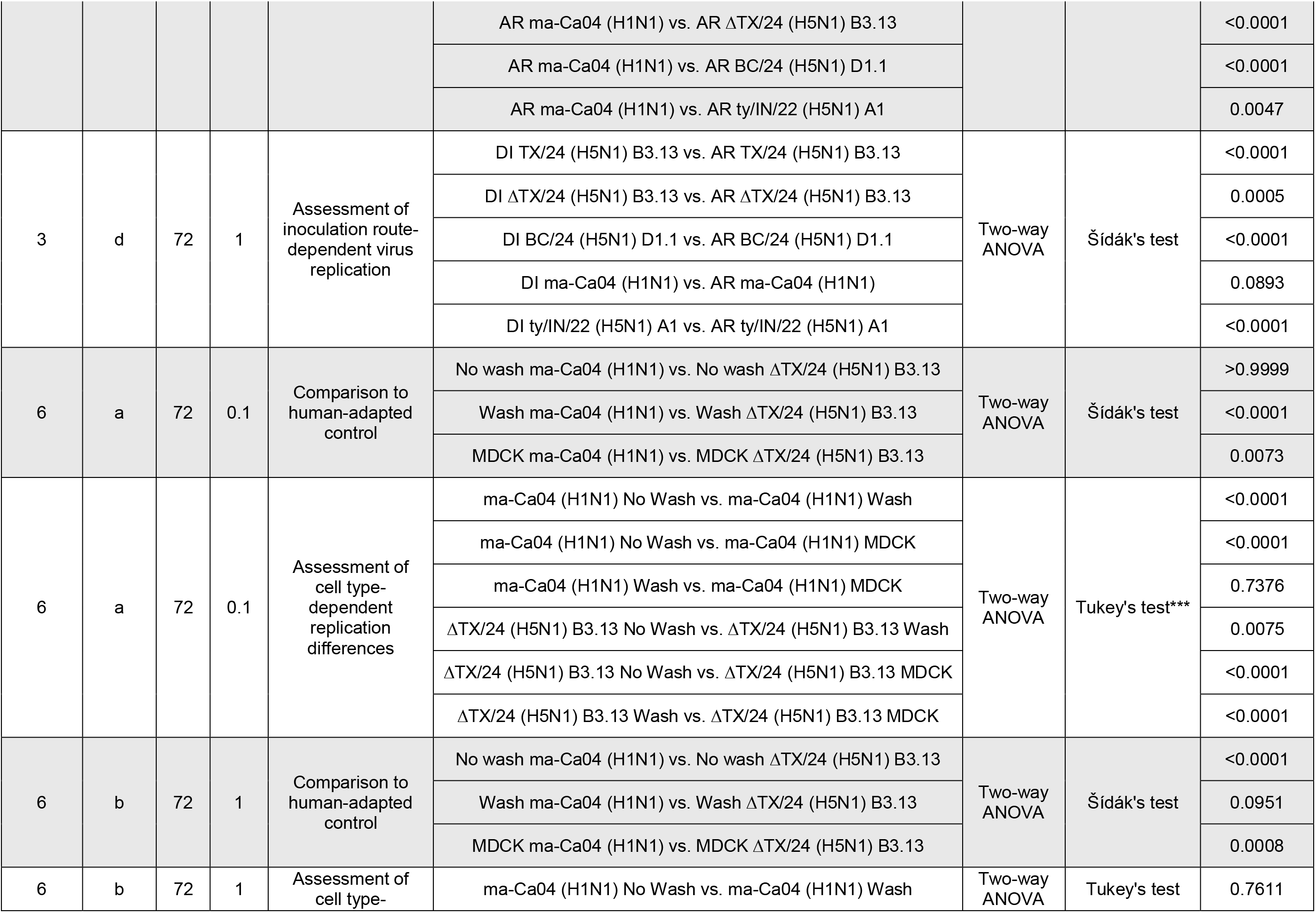
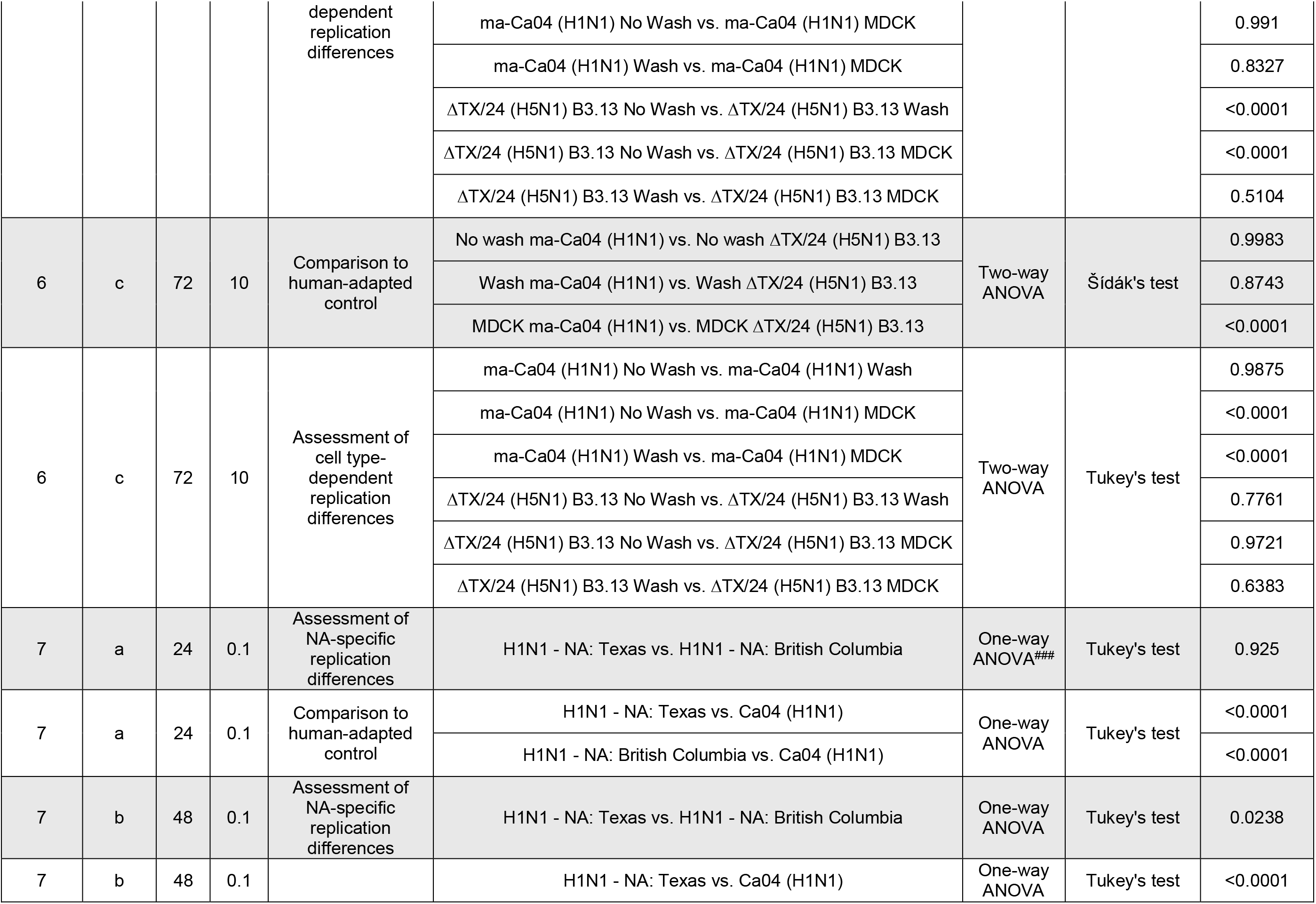
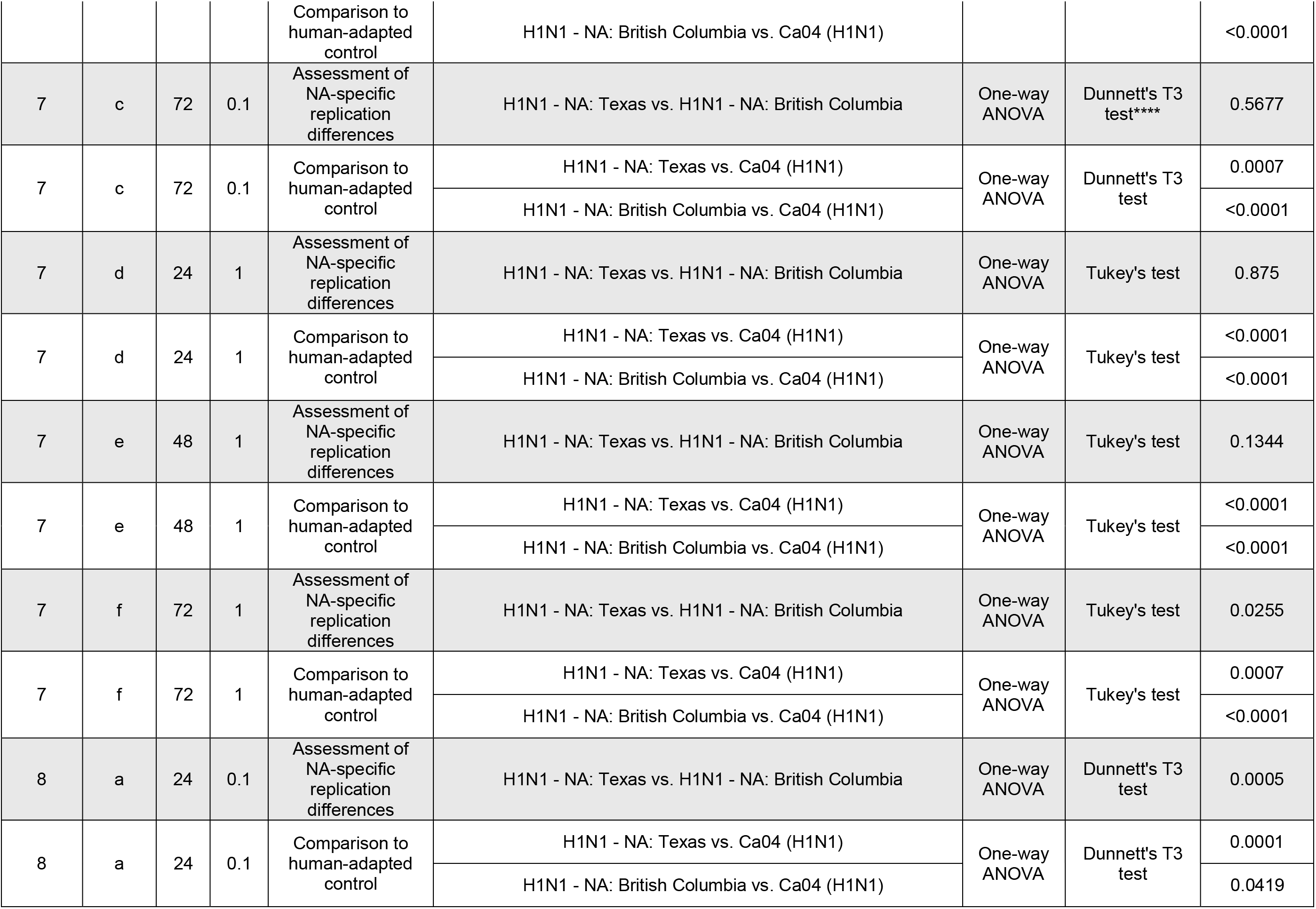
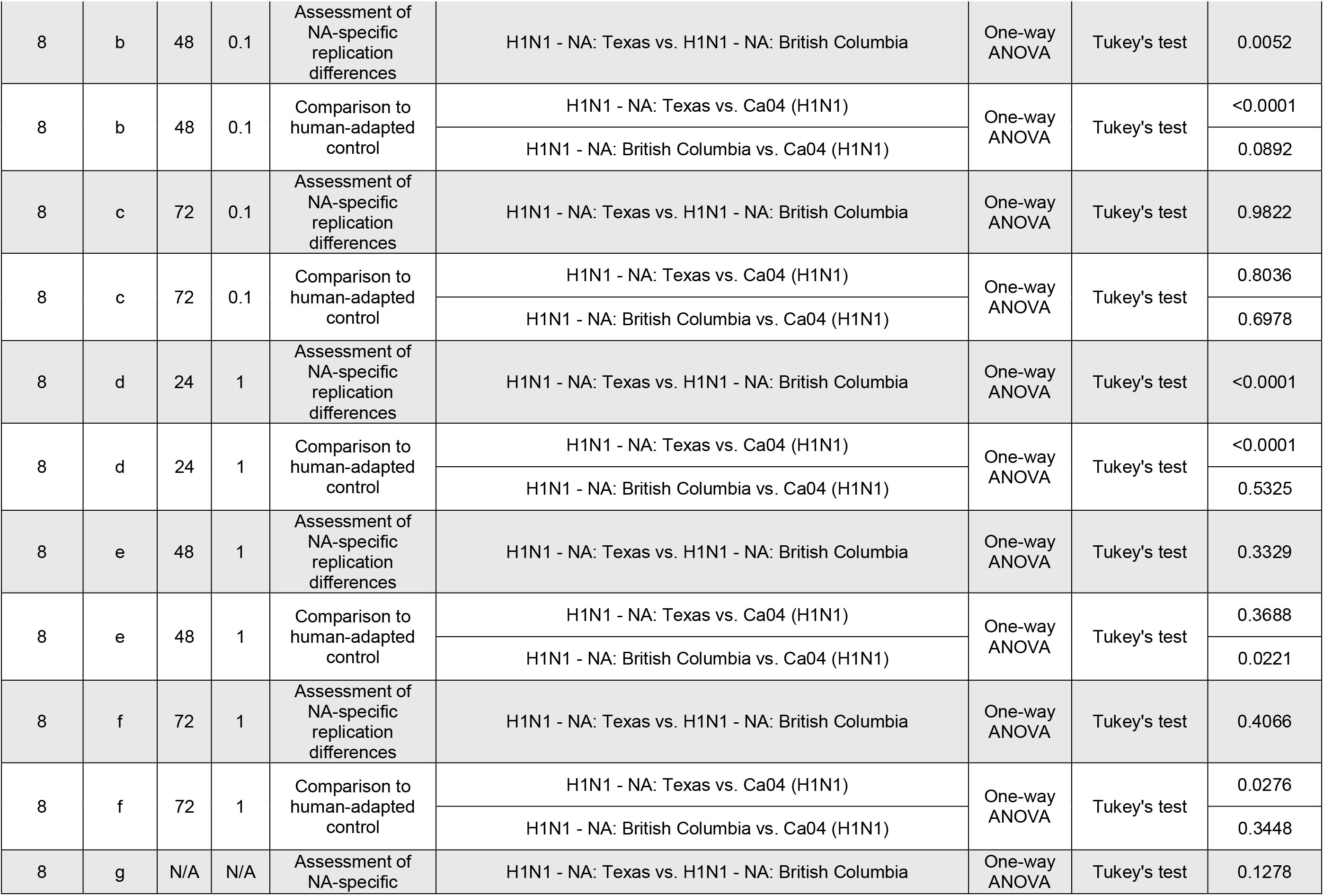
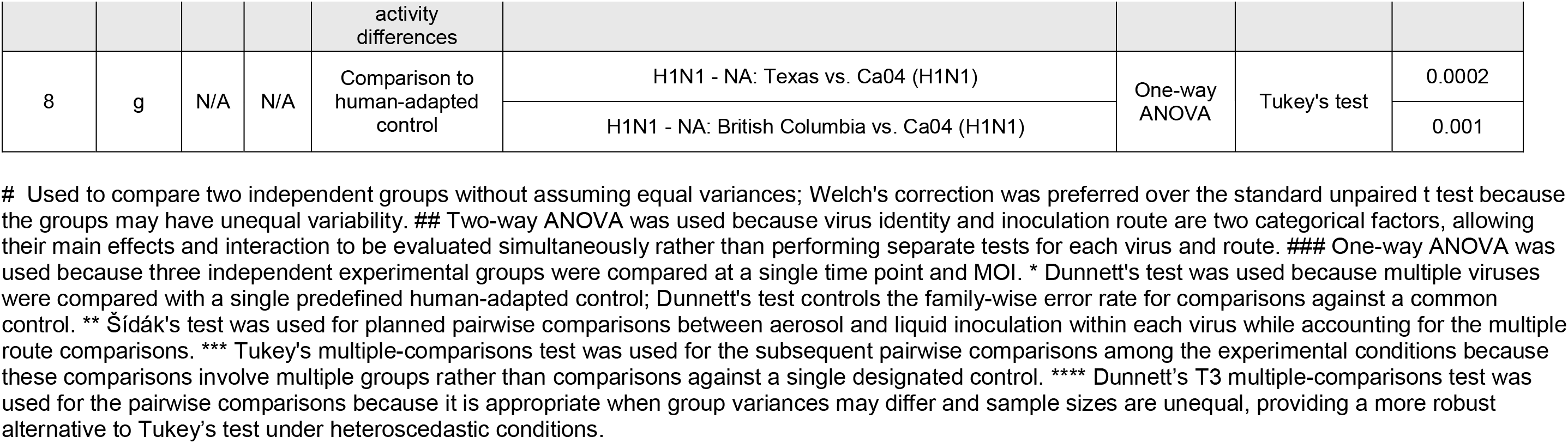
Statistical Analysis of Viral Replication, Inoculation Route Dependencies, and Neuraminidase (NA)- Specific Activity.

**SUPPLEMENTARY TABLE 3.**
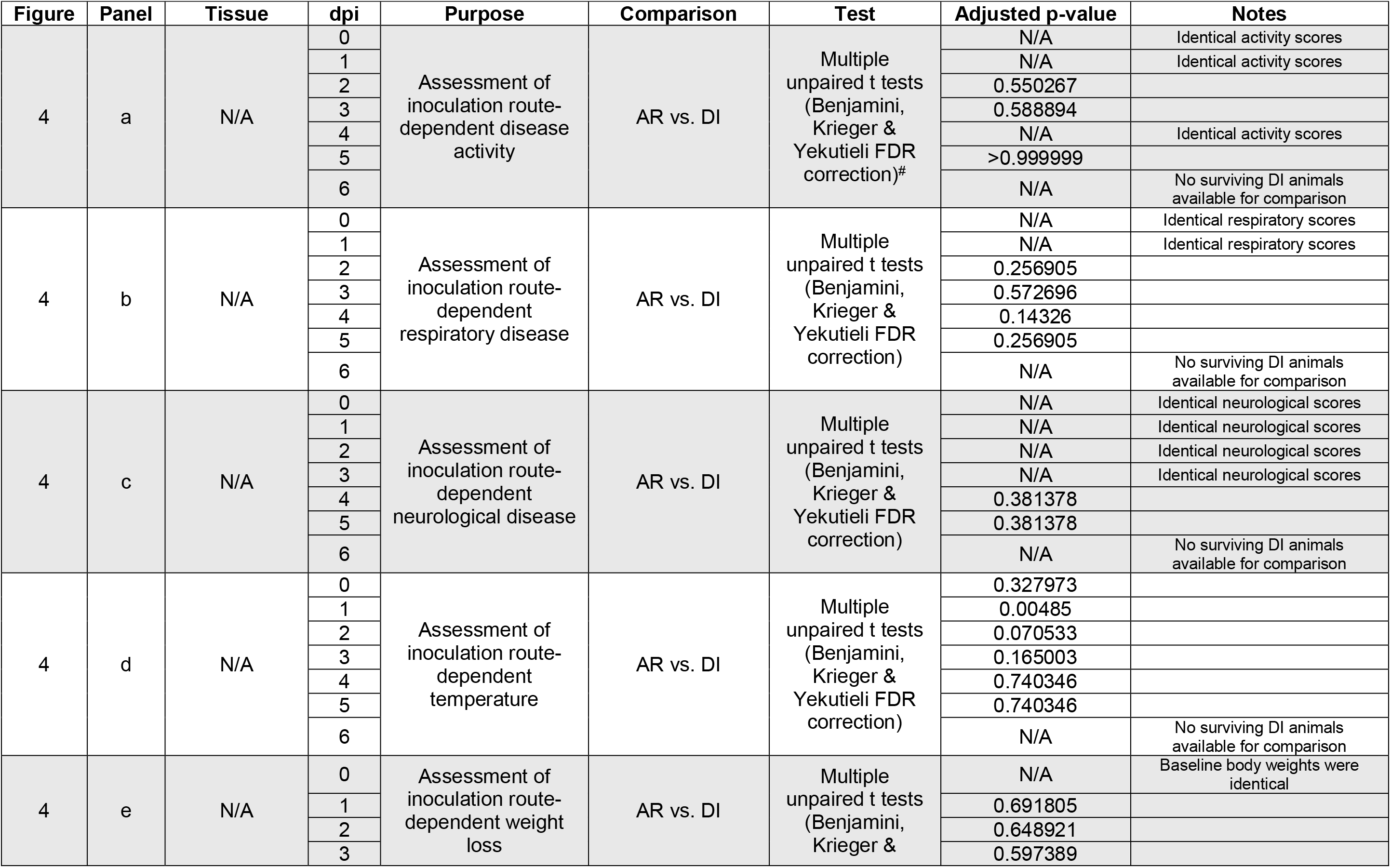
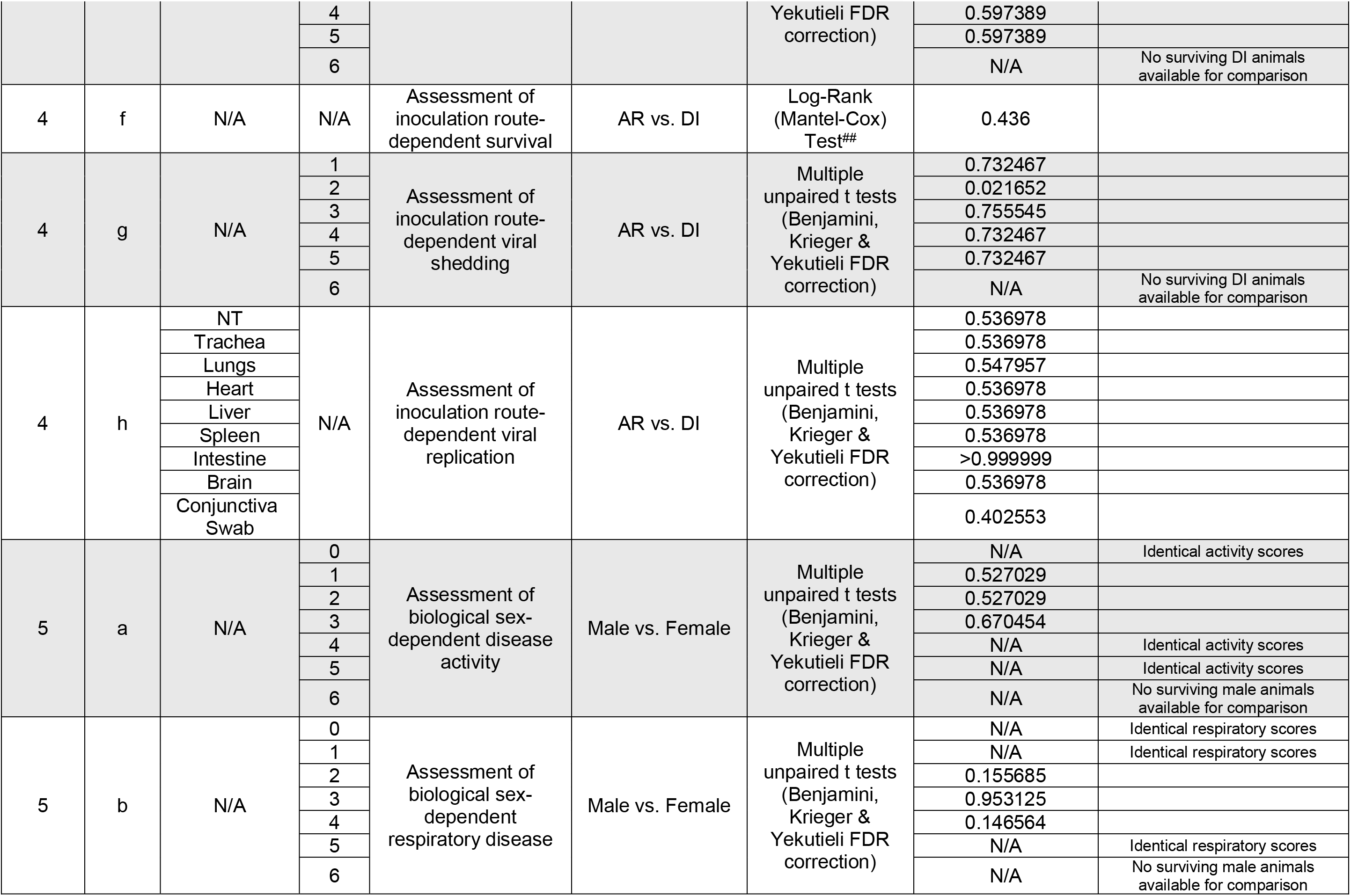
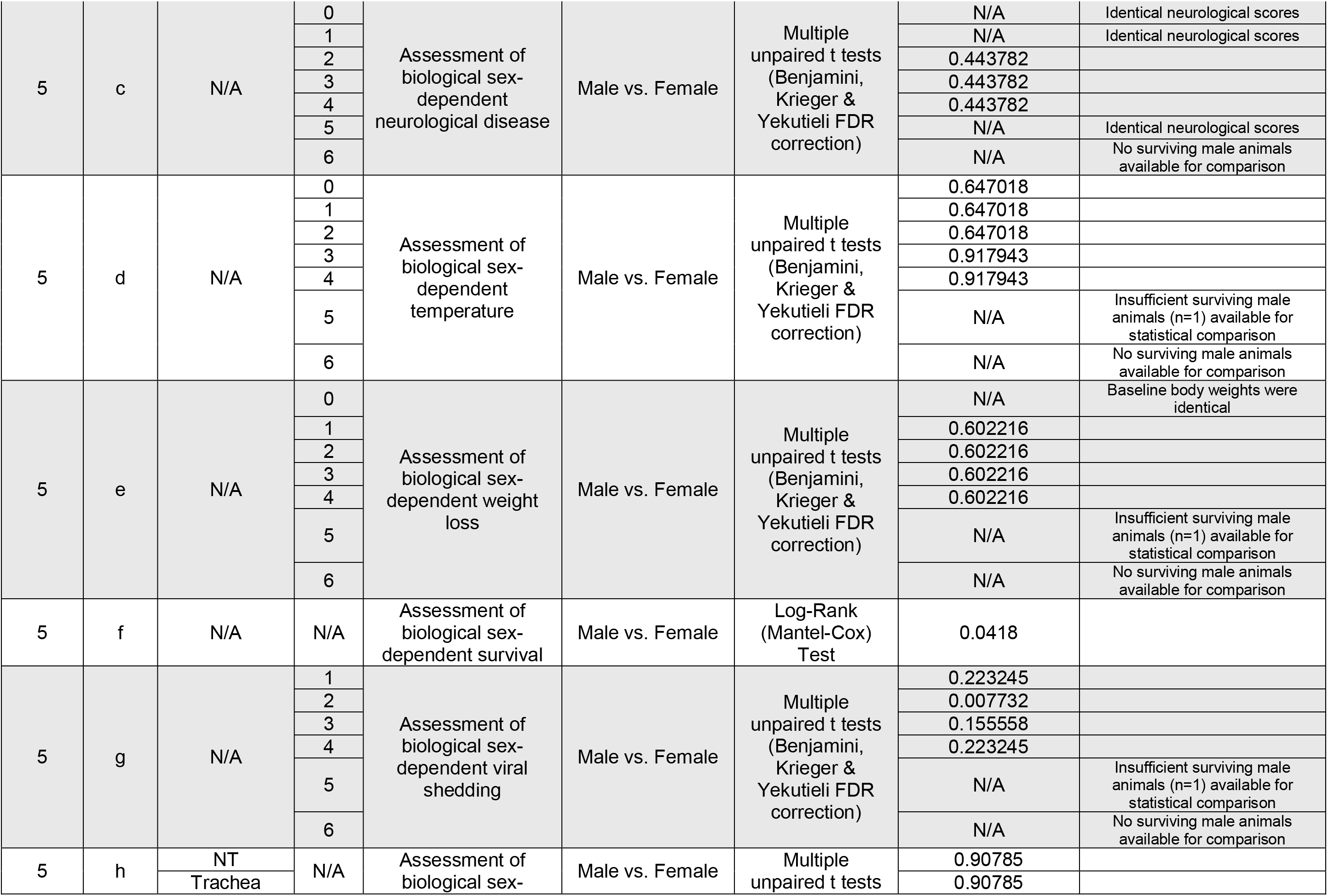
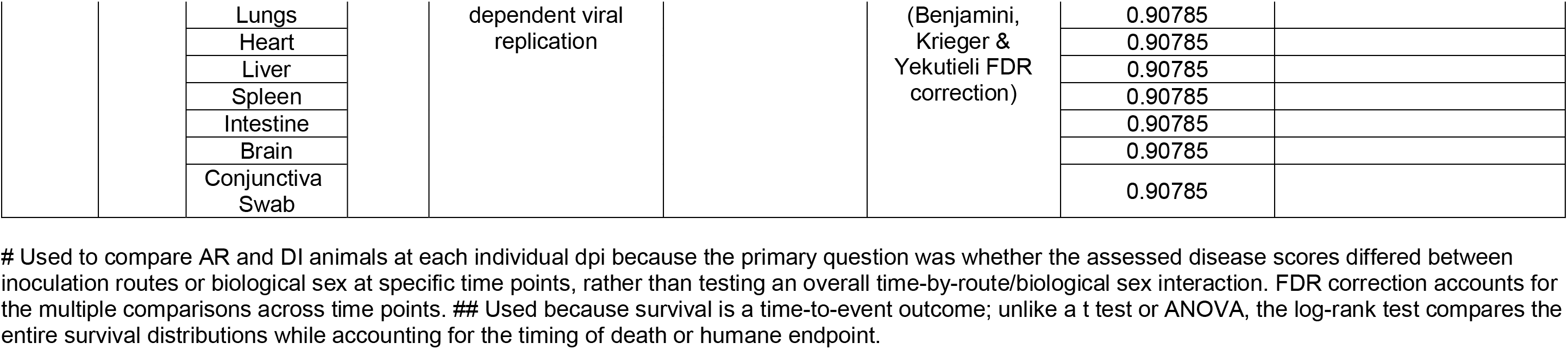
Statistical Analysis of Inoculation Route- and Biological Sex-Dependent Disease Outcomes, Survival, and Viral Shedding.

**SUPPLEMENTARY TABLE 4.**
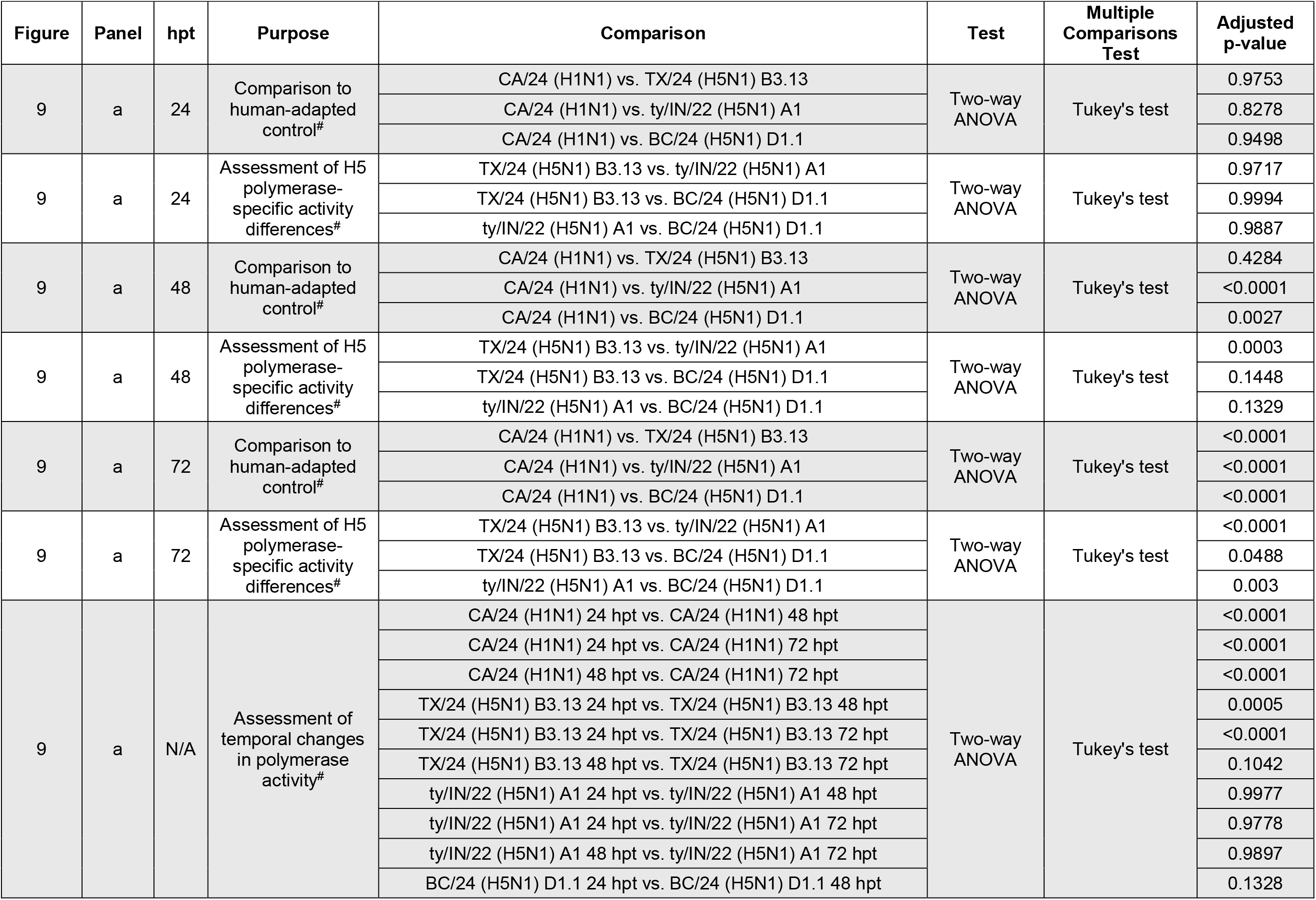
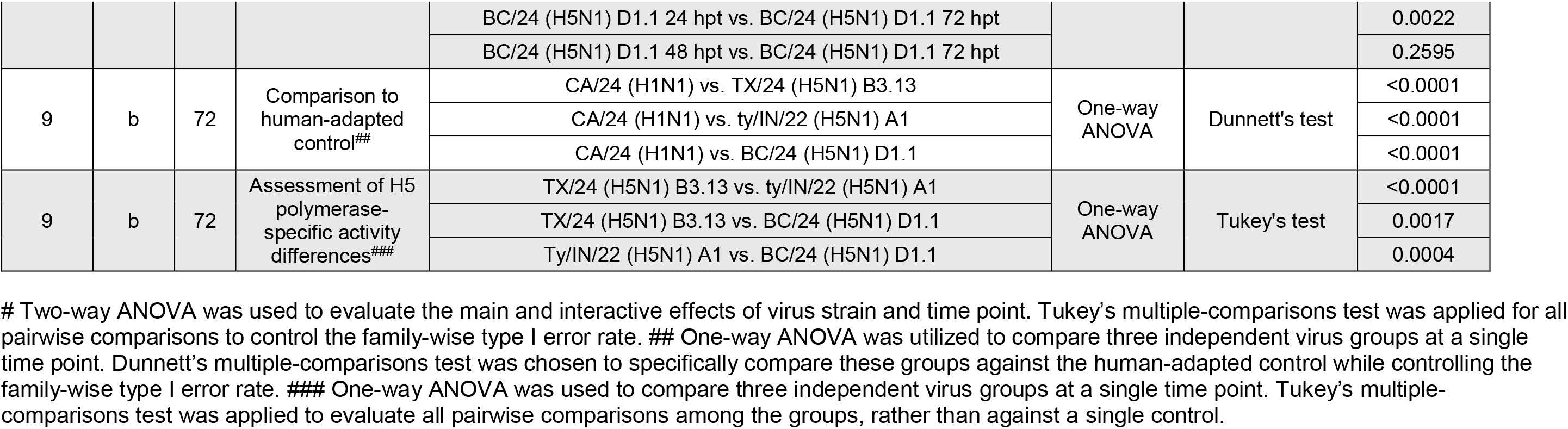
Statistical Analysis of Polymerase-Specific Activity Differences and Temporal Changes.

## REFERENCES

1. Kim JK, Negovetich NJ, Forrest HL, Webster RG. 2009. Ducks: the "Trojan horses" of H5N1 influenza. Influenza Other Respir Viruses 3:121–8.

2. Guo Y, Xu X, Wan X. 1998. [Genetic characterization of an avian influenza A (H5N1) virus isolated from a sick goose in China]. Zhonghua Shi Yan He Lin Chuang Bing Du Xue Za Zhi 12:322–5.

3. Lewis NS, Banyard AC, Whittard E, Karibayev T, Al Kafagi T, Chvala I, Byrne A, Meruyert Akberovna S, King J, Harder T, Grund C, Essen S, Reid SM, Brouwer A, Zinyakov NG, Tegzhanov A, Irza V, Pohlmann A, Beer M, Fouchier RAM, Akhmetzhan Akievich S, Brown IH. 2021. Emergence and spread of novel H5N8, H5N5 and H5N1 clade 2.3.4.4 highly pathogenic avian influenza in 2020. Emerg Microbes Infect 10:148–151.

4. Byrne AMP, James J, Mollett BC, Meyer SM, Lewis T, Czepiel M, Seekings AH, Mahmood S, Thomas SS, Ross CS, Byrne DJF, McMenamy MJ, Bailie V, Lemon K, Hansen RDE, Falchieri M, Lewis NS, Reid SM, Brown IH, Banyard AC. 2023. Investigating the Genetic Diversity of H5 Avian Influenza Viruses in the United Kingdom from 2020-2022. Microbiol Spectr 11:e0477622.

5. Turner JCM, Barman S, Feeroz MM, Hasan MK, Akhtar S, Jeevan T, Walker D, Franks J, Seiler P, Mukherjee N, Kercher L, McKenzie P, Lam T, El-Shesheny R, Webby RJ. 2021. Highly Pathogenic Avian Influenza A(H5N6) Virus Clade 2.3.4.4h in Wild Birds and Live Poultry Markets, Bangladesh. Emerg Infect Dis 27:2492–2494.

6. Chrzastek K, Lieber CM, Plemper RK. 2025. H5N1 Clade 2.3.4.4b: Evolution, Global Spread, and Host Range Expansion. Pathogens 14.

7. Koopmans MPG, Barton Behravesh C, Cunningham AA, Adisasmito WB, Almuhairi S, Bilivogui P, Bukachi SA, Casas N, Cediel Becerra N, Charron DF, Chaudhary A, Ciacci Zanella JR, Dar O, Debnath N, Dungu B, Farag E, Gao GF, Khaitsa M, Machalaba C, Mackenzie JS, Markotter W, Mettenleiter TC, Morand S, Smolenskiy V, Zhou L, Hayman DTS, One Health High-Level Expert P. 2024. The panzootic spread of highly pathogenic avian influenza H5N1 sublineage 2.3.4.4b: a critical appraisal of One Health preparedness and prevention. Lancet Infect Dis 24:e774–e781.

8. Ariyama N, Pardo-Roa C, Munoz G, Aguayo C, Avila C, Mathieu C, Almonacid LI, Medina RA, Brito B, Johow M, Neira V. 2023. Highly Pathogenic Avian Influenza A(H5N1) Clade 2.3.4.4b Virus in Wild Birds, Chile. Emerg Infect Dis 29:1842–1845.

9. Castro-Sanguinetti GR, Gonzalez-Veliz R, Callupe-Leyva A, Apaza-Chiara AP, Jara J, Silva W, Icochea E, More-Bayona JA. 2024. Highly pathogenic avian influenza virus H5N1 clade 2.3.4.4b from Peru forms a monophyletic group with Chilean isolates in South America. Sci Rep 14:3635.

10. Uhart M, Vanstreels R. 2025. Overview of high pathogenicity avian influenza H5N1 clade 2.3.4.4b in wildlife from Central and South America, October 2022-September 2025. Can J Microbiol 71:1–8.

11. Rimondi A, Vanstreets R, Nelson M, Olivera V, Gallo L, Durant A, Quintana F, Brogger M, Campana J, Dellicour S, Wolff T, Uhart MM. 2025. Persistence and spillback of mammal-adapted H5N1 genotype B3.2 viruses among South American seabirds and marine mammals. Res Sq doi:10.21203/rs.3.rs-7960151/v1.

12. Nguyen TQ, Hutter CR, Markin A, Thomas M, Lantz K, Killian ML, Janzen GM, Vijendran S, Wagle S, Inderski B, Magstadt DR, Li G, Diel DG, Frye EA, Dimitrov KM, Swinford AK, Thompson AC, Snekvik KR, Suarez DL, Lakin SM, Schwabenlander S, Ahola SC, Johnson KR, Baker AL, Robbe-Austerman S, Torchetti MK, Anderson TK. 2025. Emergence and interstate spread of highly pathogenic avian influenza A(H5N1) in dairy cattle in the United States. Science 388:eadq0900.

13. Caserta LC, Frye EA, Butt SL, Laverack M, Nooruzzaman M, Covaleda LM, Thompson AC, Koscielny MP, Cronk B, Johnson A, Kleinhenz K, Edwards EE, Gomez G, Hitchener G, Martins M, Kapczynski DR, Suarez DL, Alexander Morris ER, Hensley T, Beeby JS, Lejeune M, Swinford AK, Elvinger F, Dimitrov KM, Diel DG. 2024. Spillover of highly pathogenic avian influenza H5N1 virus to dairy cattle. Nature 634:669–676.

14. Cargnin Faccin F, Gay LC, Regmi D, Compton S, Mejias TD, Brondani JC, Joshi LR, Howerth EW, Rajao DS, Palomares RA, Perez DR. 2026. Experimental infection and viral pathogenesis of a human isolate of H5N1 highly pathogenic avian influenza strain in Jersey cows. npj Vet Sci 1:2.

15. Garg S, Reinhart K, Couture A, Kniss K, Davis CT, Kirby MK, Murray EL, Zhu S, Kraushaar V, Wadford DA, Drehoff C, Kohnen A, Owen M, Morse J, Eckel S, Goswitz J, Turabelidze G, Krager S, Unutzer A, Gonzales ER, Abdul Hamid C, Ellington S, Mellis AM, Budd A, Barnes JR, Biggerstaff M, Jhung MA, Richmond-Crum M, Burns E, Shimabukuro TT, Uyeki TM, Dugan VG, Reed C, Olsen SJ. 2025. Highly Pathogenic Avian Influenza A(H5N1) Virus Infections in Humans. N Engl J Med 392:843–854.

16. Mostafa A, Nogales A, Martinez-Sobrido L. 2025. Highly pathogenic avian influenza H5N1 in the United States: recent incursions and spillover to cattle. Npj Viruses 3:54.

17. Campbell AJ, Brizuela K, Lakdawala SS. 2025. mGem: Transmission and exposure risks of dairy cow H5N1 influenza virus. mBio 16:e0294424.

18. Harrington WN, Signore A, Kercher L, Giacinti JA, Kandeil A, Ahlstrom CA, Bevins S, Crossley B, Eure K, Fabrizio TP, Jeevan T, Lenoch J, Nolting JM, Rejmanek D, Stallknecht D, Bollinger T, Buck EJ, Carter D, Cohen BS, Dilione KE, Feddersen JC, Franks J, Goldsmith D, Highway CJ, Himsworth C, Holmes LP, Jardine C, Jahid MJ, Link P, Miller L, Nemeth NM, Owsiany M, Pybus M, Scott LC, Sharp C, Smith L, Steelman NJ, Stevens B, Berhane Y, Torchetti M, Ramey AM, Poulson R, Webby RJ. 2026. Rapid expansion of genotype D1.1 A(H5N1) influenza viruses in wild birds across North America during the 2024 migratory season. Nat Med doi:10.1038/s41591-026-04300-1.

19. Organization WH. 2024. Avian Influenza Weekly Update Number 935, February 23, 2024 ed, https://iris.who.int/server/api/core/bitstreams/39c16477-c0f6-440f-8dc5-69cff6911c26/content.

20. Organization WH. 2026. Avian Influenza Weekly Update Number 1029, January 16 2026 ed, https://cdn.who.int/media/docs/default-source/wpro---documents/emergency/surveillance/avian-influenza/ai_20260116.pdf.

21. Fabrizio TP, Kandeil A, Harrington WN, Jones JC, Jeevan T, Andreev K, Seiler P, Fogo J, Davis ML, Crumpton JC, Franks J, DeBeauchamp J, Vogel P, Daniels CS, Poulson RL, Bowman AS, Govorkova EA, Webby RJ. 2025. Genotype B3.13 influenza A(H5N1) viruses isolated from dairy cattle demonstrate high virulence in laboratory models, but retain avian virus-like properties. Nat Commun 16:6771.

22. Gu C, Maemura T, Guan L, Eisfeld AJ, Biswas A, Kiso M, Uraki R, Ito M, Trifkovic S, Wang T, Babujee L, Presler R, Jr., Dahn R, Suzuki Y, Halfmann PJ, Yamayoshi S, Neumann G, Kawaoka Y. 2024. A human isolate of bovine H5N1 is transmissible and lethal in animal models. Nature 636:711–718.

23. Eisfeld AJ, Biswas A, Guan L, Gu C, Maemura T, Trifkovic S, Wang T, Babujee L, Dahn R, Halfmann PJ, Barnhardt T, Neumann G, Suzuki Y, Thompson A, Swinford AK, Dimitrov KM, Poulsen K, Kawaoka Y. 2024. Pathogenicity and transmissibility of bovine H5N1 influenza virus. Nature 633:426–432.

24. Panova AS, Gudymo AS, Kolosova NP, Danilenko AV, Shadrinova KN, Danilchenko NV, Perfilieva ON, Moiseeva AA, Danilenko EI, Onkhonova GS, Goncharova NI, Svyatchenko SV, Vasiltsova NN, Egorova ML, Marchenko VY. 2025. Genotype A3 influenza A(H5N1) isolated from fur seals shows high virulence in mammals, but not airborne transmission. Sci Rep 15:44463.

25. Pulit-Penaloza JA, Brock N, Belser JA, Sun X, Pappas C, Kieran TJ, Basu Thakur P, Zeng H, Cui D, Frederick J, Fasce R, Tumpey TM, Maines TR. 2024. Highly pathogenic avian influenza A(H5N1) virus of clade 2.3.4.4b isolated from a human case in Chile causes fatal disease and transmits between co-housed ferrets. Emerg Microbes Infect 13:2332667.

26. Belser JA, Pulit-Penaloza JA, Brock N, Sun X, Kieran TJ, Pappas C, Zeng H, Vu MN, Lakdawala SS, Tumpey TM, Maines TR. 2025. Ocular infectivity and replication of a clade 2.3.4.4b A(H5N1) influenza virus associated with human conjunctivitis in a dairy farm worker in the USA: an in-vitro and ferret study. Lancet Microbe 6:101070.

27. Belser JA, Sun X, Pulit-Penaloza JA, Maines TR. 2024. Fatal Infection in Ferrets after Ocular Inoculation with Highly Pathogenic Avian Influenza A(H5N1) Virus. Emerg Infect Dis 30:1484–1487.

28. Kim IH, Nam JH, Kim CK, Choi YJ, Lee H, An BM, Lee NJ, Jeong H, Lee SY, Yeo SG, Lee EK, Lee YJ, Rhee JE, Lee SW, Jee Y, Kim EJ. 2024. Pathogenicity of Highly Pathogenic Avian Influenza A(H5N1) Viruses Isolated from Cats in Mice and Ferrets, South Korea, 2023. Emerg Infect Dis 30:2033–2041.

29. Pulit-Penaloza JA, Belser JA, Brock N, Kieran TJ, Sun X, Pappas C, Zeng H, Carney P, Chang J, Bradley-Ferrell B, Stevens J, De La Cruz JA, Hatta Y, Di H, Davis CT, Tumpey TM, Maines TR. 2024. Transmission of a human isolate of clade 2.3.4.4b A(H5N1) virus in ferrets. Nature 636:705–710.

30. Restori KH, Septer KM, Field CJ, Patel DR, VanInsberghe D, Raghunathan V, Lowen AC, Sutton TC. 2024. Risk assessment of a highly pathogenic H5N1 influenza virus from mink. Nat Commun 15:4112.

31. Bordes L, Gerhards NM, Roose M, Venema S, Engelsma M, van der Poel WHM, Germeraad EA, Beerens N, Vreman S. 2025. Transmission and pathogenicity in ferrets after experimental infection with HPAI clade 2.3.4.4b H5N1 viruses. J Gen Virol 106.

32. Moreira ÉA, Constant S, Constant C, Butticaz L, Wyler M, David T, Grin PM, Benarafa C, Thiel V, Alves MP, Zimmer G. 2026. Bovine-derived influenza A virus (H5N1) shows efficient replication in well-differentiated human nasal epithelial cells without requiring genetic adaptation. bioRxiv doi:10.64898/2026.01.16.699876:2026.01.16.699876.

33. Seibert B, Caceres CJ, Gay LC, Shetty N, Faccin FC, Carnaccini S, Walters MS, Marr LC, Lowen AC, Rajao DS, Perez DR. 2025. Air-liquid interface model for influenza aerosol exposure in vitro. J Virol 99:e0061925.

34. Walters MS, Gomi K, Ashbridge B, Moore MA, Arbelaez V, Heldrich J, Ding BS, Rafii S, Staudt MR, Crystal RG. 2013. Generation of a human airway epithelium derived basal cell line with multipotent differentiation capacity. Respir Res 14:135.

35. Pell TJ, Gray MB, Hopkins SJ, Kasprowicz R, Porter JD, Reeves T, Rowan WC, Singh K, Tvermosegaard KB, Yaqub N, Wayne GJ. 2021. Epithelial Barrier Integrity Profiling: Combined Approach Using Cellular Junctional Complex Imaging and Transepithelial Electrical Resistance. SLAS Discov 26:909–921.

36. Myo YPA, Camus SV, Freeberg MAT, Bernas T, Bande D, Heise RL, Thatcher TH, Sime PJ. 2025. Protocol for differentiating primary human small airway epithelial cells at the air-liquid interface. Am J Physiol Lung Cell Mol Physiol 328:L757–L771.

37. Ferreri LM, Seibert B, Caceres CJ, Patatanian K, Holmes KE, Gay LC, Cargnin Faccin F, Cardenas M, Carnaccini S, Shetty N, Rajao D, Koelle K, Marr LC, Perez DR, Lowen AC. 2025. Dispersal of influenza virus populations within the respiratory tract shapes their evolutionary potential. Proc Natl Acad Sci U S A 122:e2419985122.

38. Zanin M, Baviskar P, Webster R, Webby R. 2016. The Interaction between Respiratory Pathogens and Mucus. Cell Host Microbe 19:159–68.

39. Wagner R, Matrosovich M, Klenk HD. 2002. Functional balance between haemagglutinin and neuraminidase in influenza virus infections. Rev Med Virol 12:159–66.

40. Gaymard A, Le Briand N, Frobert E, Lina B, Escuret V. 2016. Functional balance between neuraminidase and haemagglutinin in influenza viruses. Clin Microbiol Infect 22:975–983.

41. Liu T, Wang Y, Tan TJC, Wu NC, Brooke CB. 2022. The evolutionary potential of influenza A virus hemagglutinin is highly constrained by epistatic interactions with neuraminidase. Cell Host Microbe 30:1363–1369 e4.

42. Mostafa A, Naguib MM, Nogales A, Barre RS, Stewart JP, Garcia-Sastre A, Martinez-Sobrido L. 2024. Avian influenza A (H5N1) virus in dairy cattle: origin, evolution, and cross-species transmission. mBio 15:e0254224.

43. Bauer L, Leijten L, Iervolino M, Chopra V, van Dijk L, Power M, Rijnink W, Pronk M, Spronken M, Funk M, de Vries RD, Richard M, Kuiken T, van Riel D. 2026. Attachment and replication of clade 2.3.4.4b influenza A (H5N1) viruses in human respiratory epithelium: an in-vitro study. Lancet Microbe 7:101230.

44. Fusade-Boyer M, Kocher A, Bessiere P, Figueroa T, Foret-Lucas C, Vergne T, Chevalier C, Ducatez MF, Volmer R. 2026. Investigation and impact of mammalian adaptation markers on H5N8 high pathogenicity avian influenza polymerase activity. Npj Viruses 4.

45. Dholakia V, Quantrill JL, Richardson SAS, Pankaew N, Brown MD, Yang J, Capelastegui F, Masonou T, Case KM, Ajeian J, Woodall MNJ, Magill C, Freimanis G, McCarron A, Staller E, Sheppard CM, Brown IH, Murcia PR, Smith CM, Iqbal M, Digard P, Barclay WS, Pinto RM, Peacock TP, Goldhill DH. 2026. Polymerase mutations underlie early adaptation of H5N1 influenza virus to dairy cattle and other mammals. Nat Commun 17:1603.

46. Zhang L, Lai Y, Cui Y, Yang Q, Shao Y, Ding S, Wang H, Wang L, Gao GF, Deng T. 2025. Emergence of mammalian-adaptive PB2 mutations enhances polymerase activity and pathogenicity of cattle-derived H5N1 influenza A virus. Nat Commun 17:1011.

47. Turnbull ML, Zakaria MK, Upfold NS, Bakshi S, Magill C, Das UR, Clarke AT, Mojsiejczuk L, Herder V, Dee K, Liu N, Folwarczna M, Ilia G, Furnon W, Schultz V, Chen H, Devlin R, McCowan J, Young AL, Po WW, Smollett K, Yaseen MA, Ross R, Bhide A, van Kekem B, Fouchier RAM, da Silva Filipe A, Iqbal M, Roberts E, Hughes J, Werling D, Murcia PR, Palmarini M. 2025. The potential of H5N1 viruses to adapt to bovine cells varies throughout evolution. Nat Commun 16:11042.

48. Kaftan T, Nguyen NV, Begley J, Fashina T, Carag J, Yeh S. 2025. Update on Ophthalmic Implications of Highly Pathogenic Avian Influenza A (H5N1) Virus. Pathogens 14.

49. Gadiyar I, Dobrovolny HM. 2024. Different routes of infection of H5N1 lead to changes in infecting time. Math Biosci 367:109129.

50. Kumlin U, Olofsson S, Dimock K, Arnberg N. 2008. Sialic acid tissue distribution and influenza virus tropism. Influenza Other Respir Viruses 2:147–54.

51. Thompson CI, Barclay WS, Zambon MC, Pickles RJ. 2006. Infection of human airway epithelium by human and avian strains of influenza a virus. J Virol 80:8060–8.

52. Bodewes R, Kreijtz JH, van Amerongen G, Hillaire ML, Vogelzang-van Trierum SE, Nieuwkoop NJ, van Run P, Kuiken T, Fouchier RA, Osterhaus AD, Rimmelzwaan GF. 2013. Infection of the upper respiratory tract with seasonal influenza A(H3N2) virus induces protective immunity in ferrets against infection with A(H1N1)pdm09 virus after intranasal, but not intratracheal, inoculation. J Virol 87:4293–301.

53. Ryan KA, Slack GS, Marriott AC, Kane JA, Whittaker CJ, Silman NJ, Carroll MW, Gooch KE. 2018. Cellular immune response to human influenza viruses differs between H1N1 and H3N2 subtypes in the ferret lung. PLoS One 13:e0202675.

54. Dickey BF. 2018. What it takes for a cough to expel mucus from the airway. Proc Natl Acad Sci U S A 115:12340–12342.

55. Kaler L, Iverson E, Bader S, Song D, Scull MA, Duncan GA. 2022. Influenza A virus diffusion through mucus gel networks. Commun Biol 5:249.

56. Gsell S, Loiseau E, D’Ortona U, Viallat A, Favier J. 2020. Hydrodynamic model of directional ciliary-beat organization in human airways. Sci Rep 10:8405.

57. Pohl MO, Violaki K, Liu L, Gaggioli E, Glas I, von Kempis J, Messerli C, Lin Cw, Terrettaz C, David SC, Charlton FW, Motos G, Bluvshtein N, Schaub A, Klein LK, Luo B, Leemann NC, Ammann M, Hugentobler W, Krieger UK, Peter T, Kohn T, Nenes A, Stertz S. 2025. Comparative characterization of bronchial and nasal mucus reveals key determinants of influenza A virus inhibition. mSphere 10:e0036525.

58. Corkran M, Boboltz A, Duncan GA, Scull MA. 2025. Methods for Discerning the Impact of Mucus on Host Defenses Against Viral Infection. Curr Protoc 5:e70201.

59. Kaler L, Engle EM, Corkran M, Iverson E, Boboltz A, Ignacio MA, Yeruva T, Scull MA, Duncan GA. 2025. Mucus Physically Restricts Influenza A Viral Particle Access to the Epithelium. Adv Biol (Weinh) 9:e2400329.

60. Jassem AN, Roberts A, Tyson J, Zlosnik JEA, Russell SL, Caleta JM, Eckbo EJ, Gao R, Chestley T, Grant J, Uyeki TM, Prystajecky NA, Himsworth CG, MacBain E, Ranadheera C, Li L, Hoang LMN, Bastien N, Goldfarb DM. 2025. Critical Illness in an Adolescent with Influenza A(H5N1) Virus Infection. N Engl J Med 392:927–929.

61. Steel J, Lowen AC, Mubareka S, Palese P. 2009. Transmission of influenza virus in a mammalian host is increased by PB2 amino acids 627K or 627E/701N. PLoS Pathog 5:e1000252.

62. Fan S, Hatta M, Kim JH, Halfmann P, Imai M, Macken CA, Le MQ, Nguyen T, Neumann G, Kawaoka Y. 2014. Novel residues in avian influenza virus PB2 protein affect virulence in mammalian hosts. Nat Commun 5:5021.

63. Kim YI, Jang SG, Kwon W, Kim J, Park D, Choi I, Choi JH, Gil J, Yu M, Jeong B, Kim EH, Kim SM, Kim H, Ahn JW, Hwang S, Heo SY, Casel MAB, Rollon R, Fabrizio T, Webby RJ, Choi YK. 2025. PB2 and NP of North American H5N1 virus drive immune cell replication and systemic infections. Sci Adv 11:eady1208.

64. Ye J, Sorrell EM, Cai Y, Shao H, Xu K, Pena L, Hickman D, Song H, Angel M, Medina RA, Manicassamy B, Garcia-Sastre A, Perez DR. 2010. Variations in the hemagglutinin of the 2009 H1N1 pandemic virus: potential for strains with altered virulence phenotype? PLoS Pathog 6:e1001145.

65. Caceres CJ, Gay LC, Faccin FC, Regmi D, Palomares R, Perez DR. 2024. Influenza A(H5N1) Virus Resilience in Milk after Thermal Inactivation. Emerg Infect Dis 30:2426–2429.

66. Reed LJ, Muench H. 1938. A SIMPLE METHOD OF ESTIMATING FIFTY PER CENT ENDPOINTS. American Journal of Epidemiology 27:493–497.

67. Pena L, Vincent AL, Ye J, Ciacci-Zanella JR, Angel M, Lorusso A, Gauger PC, Janke BH, Loving CL, Perez DR. 2011. Modifications in the polymerase genes of a swine-like triple-reassortant influenza virus to generate live attenuated vaccines against 2009 pandemic H1N1 viruses. J Virol 85:456–69.

68. Santos JJS, Abente EJ, Obadan AO, Thompson AJ, Ferreri L, Geiger G, Gonzalez- Reiche AS, Lewis NS, Burke DF, Rajao DS, Paulson JC, Vincent AL, Perez DR. 2019. Plasticity of Amino Acid Residue 145 Near the Receptor Binding Site of H3 Swine Influenza A Viruses and Its Impact on Receptor Binding and Antibody Recognition. J Virol 93.

